# Clair3-Trio: high-performance Nanopore long-read variant calling in family trios with Trio-to-Trio deep neural networks

**DOI:** 10.1101/2022.05.03.490460

**Authors:** Junhao Su, Zhenxian Zheng, Syed Shakeel Ahmed, Tak-Wah Lam, Ruibang Luo

## Abstract

Accurate identification of genetic variants from family child-mother-father trio sequencing data is important in genomics. However, state-of-the-art approaches treat variant calling from trios as three independent tasks, which limits their calling accuracy for Nanopore long-read sequencing data. For better trio variant calling, we introduce Clair3-Trio, the first variant caller tailored for family trio data from Nanopore long-reads. Clair3-Trio employs a Trio-to-Trio deep neural network model, which allows it to input the trio sequencing information and output all of the trio’s predicted variants within a single model to improve variant calling. We also present MCVLoss, a novel loss function tailor-made for variant calling in trios, leveraging the explicit encoding of the Mendelian inheritance. Clair3-Trio showed comprehensive improvement in experiments. It predicted far fewer Mendelian inheritance violation variations than current state-of-the-art methods. We also demonstrated that our Trio-to-Trio model is more accurate than competing architectures. Clair3-Trio is accessible as a free, open-source project at https://github.com/HKU-BAL/Clair3-Trio.

## Introduction

Accurate identification of genetic variants in family trios in the human genome is an important task in genomics, which provides insight into precision medicine and phenotype understanding [1]. The human genome follows the Mendelian inheritance [2], with half of the child’s genome in family trios inherited from each parent. Calling genetic variants in trios provides a more comprehensive understanding of the inheritance pattern of genetic variants in families [3].

Several state-of-the-art deep learning-based methods are available for calling small variants from Oxford Nanopore Technologies (ONT) data. They are based on two main designs: pileup and full-alignment. Clairvoyante [4], Clair [5] and Nanocaller [6] use a pileup-based design, which summarizes the read alignments into features and counts, which are then piped into a variant-calling network. PEPPER-Margin-DeepVariant (PEPPER) [7], on the other hand, applies a haplotype-aware variant calling pipeline and uses full alignment-based input to call variants via neural networks. Clair3 [8] combines the two major designs, using an advance and cascade design, which symphonizes pileup for the best speed and full-alignment for the best accuracy for calling variants from ONT data. Other variant-calling methods, including Medaka [9] and Longshot [10], are also available for ONT data. However, all the state-of-the-art methods are designed for calling individual variants from trios and fail to leverage Mendelian inheritance in the family for better variant-calling accuracy for ONT data.

For calling varaints with genetic information shared in family trios, two pilot studies based on DeepVariant [11] have been developed. dv-trio [12] provides a processing pipeline to call variants using DeepVariant, together with GATK [13] and FamSeq [14], to reduce the number of Mendelian inheritance violations in its variant calling. DeepTrio [15] extends DeepVariant’s single sample input to accept the input of three samples in its deep neural networks to call candidate sites identified by heuristic checking. Current trio variant callers do not include Mendelian inheritance violation factors in their model architecture designs or decisions. Furthermore, all these methods are designed for Illumina and PacBio HiFi data, and cannot call variants from ONT data. Therefore, there is currently no trio information-aware caller available for calling variants from ONT data.

Generally, two research gaps remain for calling variants from trios for ONT data: (1) how to train the model to learn from the information about both individuals and that preserved in family trios; and (2) how to train the model to predict following Mendelian inheritance, a basic feature in family trios. Unfortunately, these two questions have never been studied in the ONT data and remain unsolved in the community.

To fill the two main research gaps and improve variant calling from trios’ ONT data, we propose a new model: Clair3-Trio. Clair3-Trio is the first variant caller tailored for family trios ONT data with a Trio-to-Trio deep neural network model design that allows it to input the trio’s sequencing information and output all of the trio’s predicted variants. Using the Trio-to-Trio model, Clair3-Trio can efficiently call variants based on individual and family trio information. We also designed a loss function, **MCVLoss** (**M**endelian Inheritance **C**onstraint **V**iolation **Loss**), to make the model explicitly encode the priors of Mendelian inheritance in trios to improve its variant calling (described in the **Methods** section). Based on our experiment on the Genome in a Bottle (GIAB) HG002 trio data [16], Clair3-Trio showed comprehensive improvement in experiments compared to state-of-the-art methods. It showed an increment of over +10% in the F1-score of the child and +5% in the F1-score of the parents compared to Clair3 and PEPPER when tested at 10x ONT data. In addition, it showed an order of magnitude fewer Mendelian inheritance violations than other methods. All codes and experimental settings for Clair3-Trio are publicly available at https://github.com/HKU-BAL/Clair3-Trio.

## Methods

### Family trio variant calling with Clair3-Trio

Clair3-Trio consists of two main modules (**Figure 1A**): (1) data preprocessing, which uses the Clair3 pileup model and WhatsHap phase, as well as the haplotag sub-module [17] function to phase the data of each individual in a family; and (2) model calling, which calls family trio variants with the Clair3-Trio model. The inputs for Clair3-Trio are three alignment files from a family trio: child, mother and father. The workflow and model are discussed in the following.

**Figure 1.**
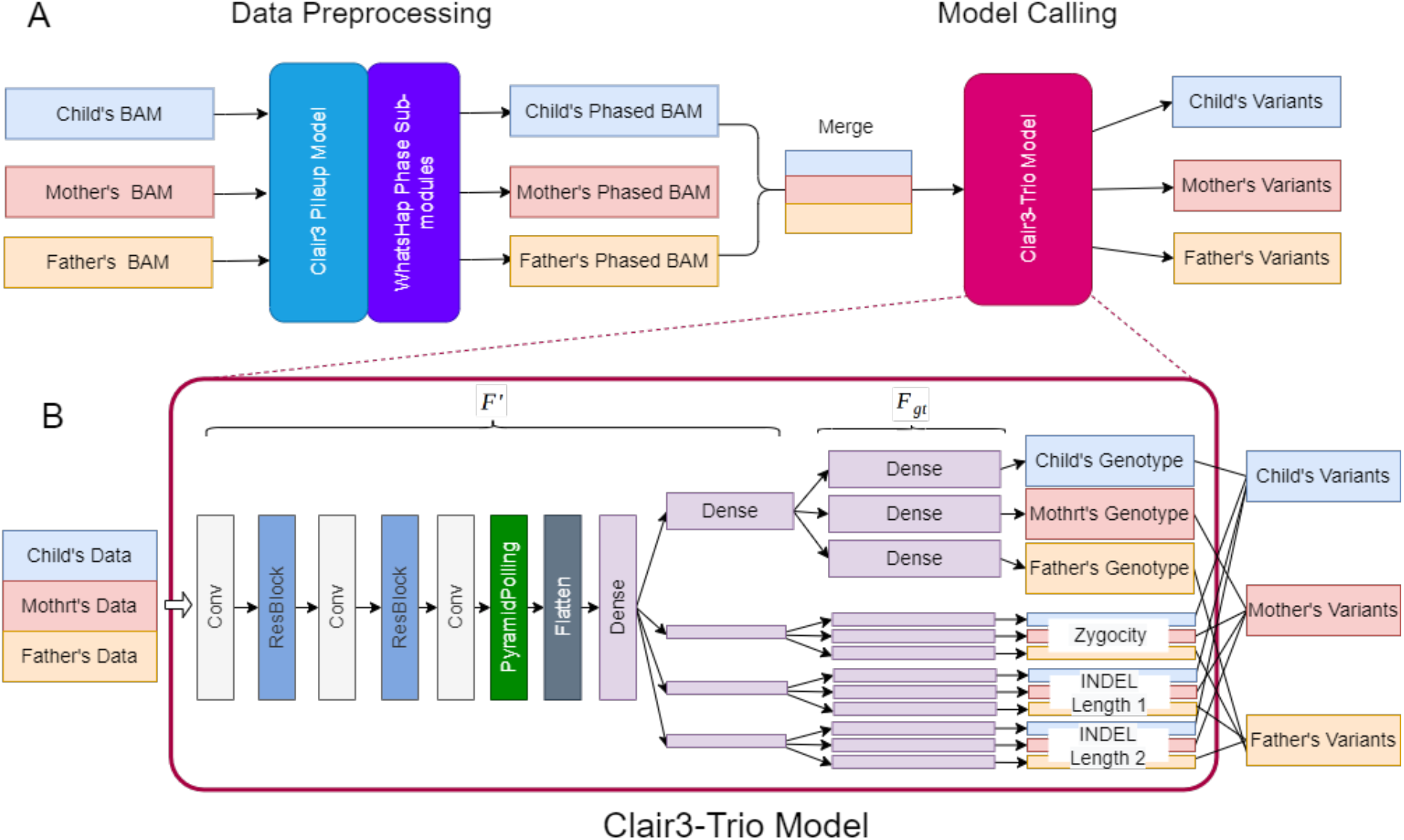
Workflow of Clair3-Trio. **A.** The calling workflow of Clair3-Trio. The trio’s sequencing data are first phased by Clair3’s pileup variant calling and WhatsHap phase submodules and then fed into the Clair3-Trio model to call variants in the trio. **B.** The architecture of the Clair3-Trio model.

#### Data preprocessing

For data preprocessing, first, we use the Clair3 pileup model to efficiently find all genetic variants that can be easily predicted with high confidence, and then we use WhatsHap to obtain all phase variants and haptag reads, based on the called heterozygous single nucleotide polymorphisms (SNP) to get phased alignments for the Clair3-Trio model. With all individuals’ haptaged alignments available, we use a simple heuristic approach to identify candidate positions that might have any genetic variants in the family, as follows: (1) the Clair3 pileup model grasps all positions with supporting alternative allele frequency exceeding 0.08 and outputs all individual variant and non-variant calling with confidence scores [8]. (2) Next, all pileup variants called and 20% of low-quality pileup reference calls are collected from each individual as the individual’s potential variant candidate sites. (3) Then we unite all the potential variants of each individual in the family as the trio’s variant candidates. Thus, any variants identified in a sample can be treated as candidates in the Clair3-Trio model.

#### Clair3-Trio model: a Trio-to-Trio deep neural network model

The Clair3-Trio model is a Trio-to-Trio model that can input all alignments from the family trio and output all variants from the same trio. The inputs for the Clair3-Trio model are generated by merging phased full alignments from trios. For each individual, the full-alignment information is converted into eight different feature channels, as previously discussed for Clair3 [8]. For each channel, we aggregate the same channel from each individual in the same family order as the input of the Clair3-Trio model (**Figure 1B**).

The neural network of the Clair3-Trio model consists of multiple layers: convolutional layers (Conv), residual convolutional layers (ResBlock), pyramid pooling layers, and dense layers (**Figure 1B)**. Clair3-Trio uses independent dense layers to predict each individual’s genotype, zygosity, and two insertion or deletion (INDEL) lengths in the last layer. All outputs from the model are then combined and converted to variant records for each individual.

#### Training a Clair3-Trio model

To train a Clair3-Trio model in family trio data, we applied (1) a label cleaning module (Representation Unification) to clean the training data, and (2) a trio data filtering module (MCV filtering) to further filter Mendelian violation sites in the training data. The two modules were established based on experiments. We use the Representation Unification module from Clair3 to unify the true variants label with the alignment information in the training data. The Representation Unification model may include Mendelian conflict in the unification process. We added MCV filtering to discard a few candidate sites (0.05% of candidate sites) in training data that violated Mendelian inheritance constraints. After cleaning the data, we performed random downsampling to make the model increase its generalization at different levels of data coverage. We downsampled the data into a range of coverage of 10x, 30x, 60x, and 80x for all samples, kept the child data at high coverage, and downsampled only the parent samples for low coverage. After downsampling, we kept 30% of the data of each coverage combination to balance speed and performance, leading to 33,353,000 candidates (from the GIAB HG002 family) in our training dataset. With the training dataset available, Clair3-Trio was trained in a two-step procedure. First, we trained an initial model of Clair3-Trio via the focus loss function, and then we fine-tuned the initial Clair3-Trio model with the addition of multiple task MCVLoss function. We also tried other training techniques, but they failed to improve Clair3-Trio. This is elaborated in the **Supplementary Notes**.

#### Differences between Clair3-Trio and the Clair3 full-alignment model

Our approach differs from Clair3 mainly in the following ways:

1. Targeted for best accuracy, Clair3-Trio can call variants in all potential variant sites in a family, while Clair3 calls them in individuals. Clair3-Trio has much more relaxed candidate selection criteria for variant candidate selection than Clair3. Claire3-Trio has 100% of variants and 20% of reference sites, compared to 30% and 10%, respectively, in Clair3, so Clair3-Trio ends up with 2.2 times more candidates. On the other hand, the variant candidates in the Clair3-Trio model are the union of all trio members, resulting in 1.9 times more candidates than individual variants. Typically, Clair3-Trio calls 4.2 times more candidates, on average, than the default Clair3 for each sample.
2. Clair3-Trio is a Trio-to-Trio model, which uses all data in the trio to predict all of the trio members’ variants directly and consistently, while Clair3 is a powerful individual variant caller, which can be treated as a One-to-One model. More information about the Trio-to-Trio and the One-to-One model is provided in the **Results** section.
3. The Clair3-Trio model uses the MCVLoss function for fine-tuning, adding penalties to the trio’s variant predictions that violate the Mendelian constraints, giving Clair3-Trio a comprehensive understanding of the family trio’s variant calling.

With the Trio-to-Trio model’s architecture and MCVLoss function, our model is well-tailored for calling variants in a family, resulting in a substantial improvement in all the benchmark experiments (see the **Results** section) using the same training data as in Clair3.

### Modeling Mendelian inheritance with MCVLoss in deep neural networks

The Trio-to-Trio model can predict the trio’s variants with trio’s information, but how to explicitly add the Mendelian inheritance information to the model remains an open question. In the following subsections, we discuss the MCVLoss function, which is designed to control the Mendelian inheritance violation rate in the model. We briefly describe the original loss function in Clair3 and then introduce the MCVLoss function.

#### Loss function for a single sample

First, we detail the original loss function for an individual, inherited from Clair3, to better illustrate the basic components in the Clair3-Trio loss function. The output of Clair3 includes four variant tasks – genotype, zygosity and two INDEL length tasks – as previously described in Clair3 [8]. The most important task in Clair3 is to predict the genotypes, which are classified into 21 genotypes. If X denotes the alignment from a single sample, the probability of each possible genotype from the 21 genotypes for each sample is:

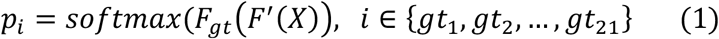

where *F_gt_* represents the Clair3 model’s last layer – the 21-genotype outputting layer – and *F′* represents all the other Clair3 layers, other than the last *F_gt_* layer, as in **Figure 1B**. Based on the probability of 21 genotypes, the loss function of Clair3 can be simplified as:

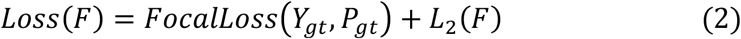

where *Y_gt_* denotes the true 21-genotype label, *P_gt_* denotes the predicted probability of each 21-genotype label, and *L*_2_ denotes the L2 regularization terms of the model. We ignore the zygosity and INDEL length terms in this simplified formula for simplicity (their formulas are identical to the 21 genotypes task). For applications, the complete loss functions, including 21 genotypes, zygosity, INDEL length 1 and INDEL length 2, are described in the Clair3 paper [8].

#### The output of Clair3-Trio and the computation of the trio probability

We extended the model output in Clair3 from the individual to compute trio genotypes in Clair3-Trio. The probability of the trio members is represented as:

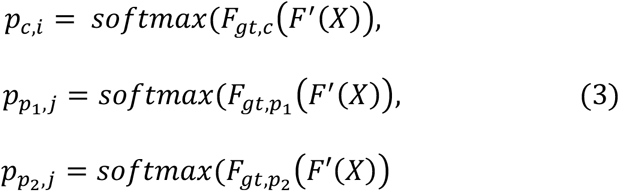

where the *F′* represents all layers of Clair3-Trio except for the last layer, and *F_gt,c_*, *F*_*gt,p*_1__, *F*_*gt,p*_2__ represents the last three fully connected layers for computing the 21 corresponding child, parent-1 and parent-2 genotypes. Parent-1 can be the mother or father in the trio, and parent-2 is the remaining parent. The probability of each trio genotype in the family is computed as:

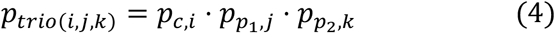

For each individual’s probability, we simply have the property that:

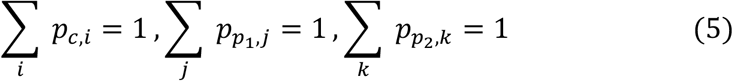

Combining formulas (4) and (5) for the trio genotype, we have a similar property for the trio’s probability:

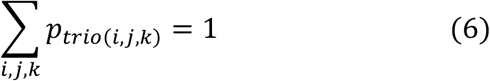

#### The Mendelian constraint violation loss function: MCVLoss

MCVLoss is based on the idea of penalizing the trio genotype that violates the Mendelian inheritance. For each trio genotype, we define a parameter *β*, representing the valid degree of the genotype:

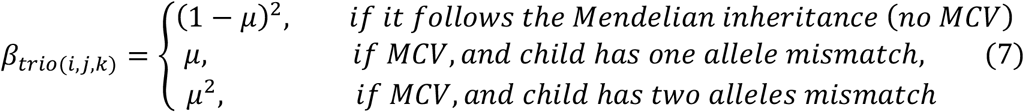

where *μ* is the mutation rate per generation, set as 1e-8 by default [18]. Combining the probability of each trio genotype in the family and the corresponding valid degree, the predicted overall valid degree for trio prediction becomes:

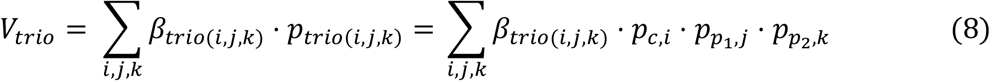

Based on formulas (6), (7) and (8), we know that *V_trio_* ∈ (0,1). With all this information, the MCVLoss is defined as:

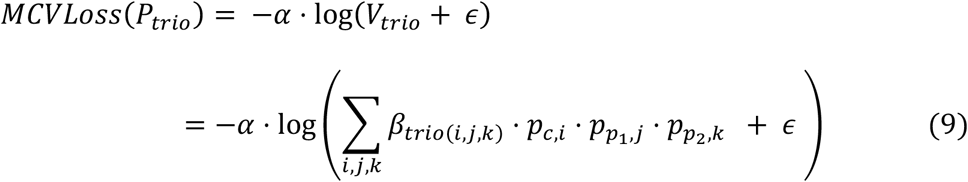

where *α* controls the importance of the Mendelian inheritance penalty in the model, and *ϵ* is a small number (1e-9 by default) to cap the log function to avoid reaching infinity. *α* is set as 1 by default, which was decided experimentally.

With the MCVLoss available, the final Clair3-trio loss function is:

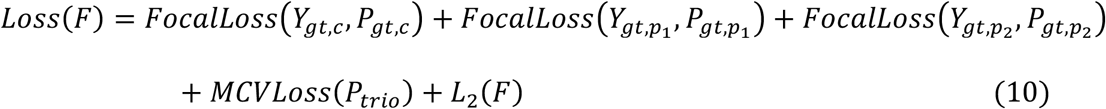

where *Y_gt_* denotes the true 21-genotype ground truth and *P_gt_* denotes the predicted probability of each 21-genotype label.

In this manner, MCVLoss introduced the Mendelian inheritance prior to model training. The detailed results of using MCVLoss are presented in the **Results** section.

### Benchmarking methods and metrics

We use Precision, Recall, and F1-score metrics to evaluate the family trio variant-calling performance in different configurations. The Precision, Recall and F1-score are computed via hap.py (v0.3.12) [19]. We computed the number of Mendelian violation variants in trios using the following steps: (1) merging all trio variants results using BCFtools (v1.12) [20] with the flag “-f PASS -0 -m all”, and (2) computing the number of Mendelian violations via RTG tools (v3.12.1) [21]. We also computed the number of *de novo* variants in the model’s prediction, where the *de novo* variants [15] are defined as variants confidently genotyped as 0/1 in the child and as 0/0 or unknown in the parents. Note that the metrics of Precision, Recall, F1-score, and number of *de novo* variants are constrained in the confidence region, while the number of the Mendelian violations is computed in all sites.

## Results

### Data description

We conducted our experiments on the dataset collected in the Genome in a Bottle (GIAB) [16] Ashkenazi Jewish trio (HG002-child, HG003-father, HG004-mother). We obtained the ONT sequencing data from the Human Pangenome Reference Consortium (HPRC) [22], with high coverage in three samples, HG002 (^~^432x), HG003 (^~^85x), and HG004 (^~^88x), which were base-called via Guppy4.2.2. We trained models on the ONT data while holding out chromosome 20 in all training stages and preserving it for testing. The truth variants for the trio were obtained from GIAB’s v4.2.1 small variant benchmark [16]. We compared Clair3-Trio with Clair3 (v0.1-r6) and PEPPER-Margin-DeepVariant (r0.4) (PEPPER). For individual evaluation, the benchmark was constrained in the individual region provided in GIAB’s v4.2.1 small variant benchmark, while the computation of *de novo* variants was constrained in the trio’s overlapped high-confident bed regions.

### Assessing variant-calling accuracy in individuals

We compared the Clair3-Trio variant-calling performance against Clair3 and PEPPER at different coverage in individuals from the GIAB trio. The overall benchmark results are shown in **Figure 2** (SNP+INDEL), with SNP and INDEL breakdowns in **Supplementary Figure 1** (SNP) and **Supplementary Figure 2** (INDEL). For all variants, we observed that Clair3-Trio had a better performance in the F1-score than Clair3 and PEPPER. The performance gain was especially profound in the lower coverage data. Clair3-Trio achieved an F1-score of 92.85% and 92.12% at 10x data from HG002 (child) and HG003 (parent), compared to 82.78% and 86.77% in Clair3, and 54.77% and 65.70% in PEPPER, respectively. Clair3-Trio was significantly better than Clair3 (*p*-value < 0.01, two-tailed t-test) and PEPPER (*p*-value < 0.001, two-tailed t-test) at 10x coverage. In high coverage data, at 70x to 90x, the Clair3-Trio performance improvement over Clair3 and PEPPER was less significant. More details are provided in **Supplementary Table 1**.

**Figure 2.**
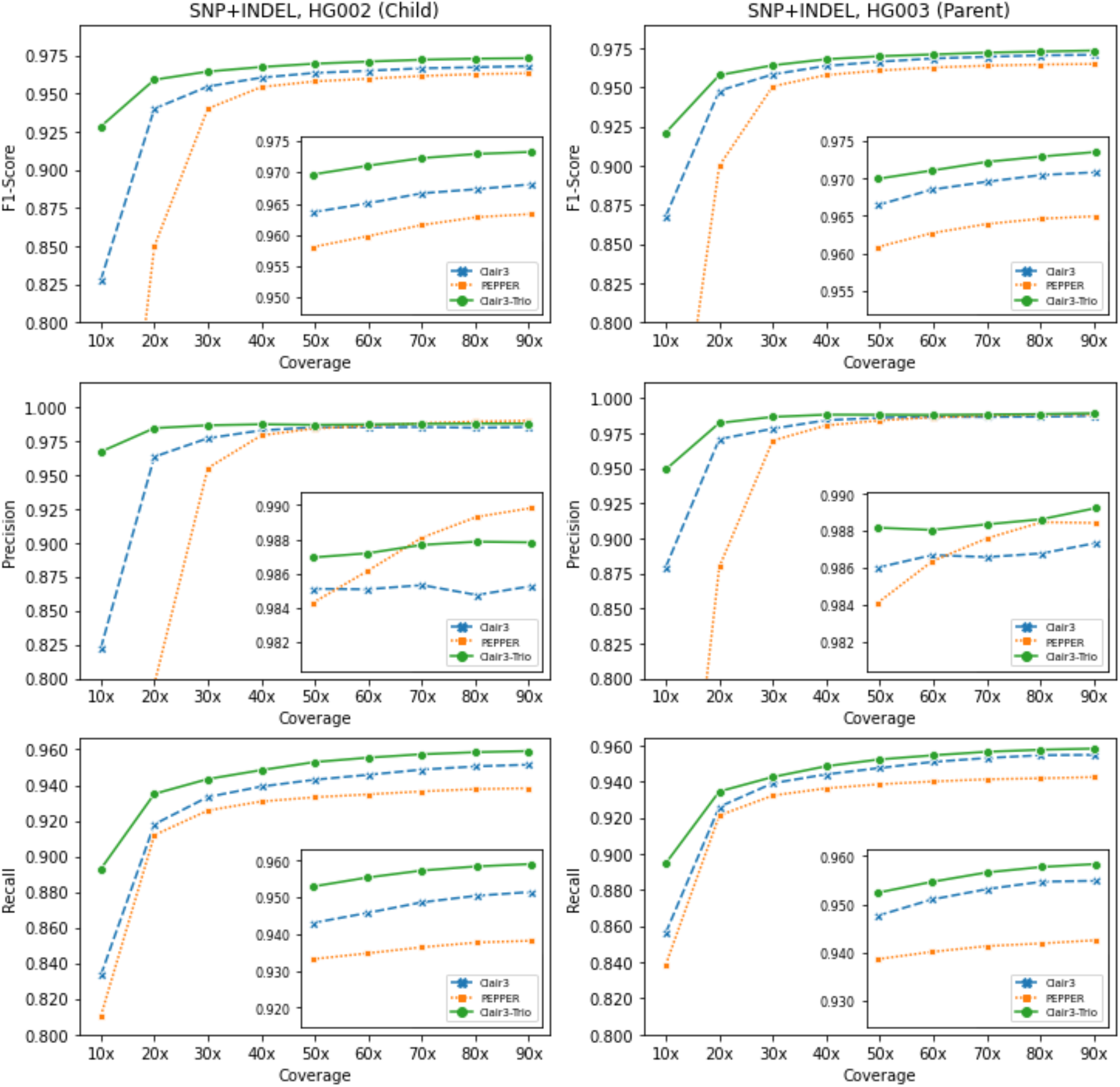
Overall benchmarking results on the GIAB trio. The SNP+INDEL’s F1-score, Precision and Recall of different tools at coverage from 10x to 90x on HG002 (child, left) and HG003 (parent, right).

The SNP and INDEL performance breakdowns are shown in **Supplementary Figure 1** and **2**. We found that Clair3-Trio had higher performance in both SNP and INDEL than Clair3 and PEPPER. For SNP, Clair3-Trio performed better than Clair3 and PEPPER, especially below 40x coverage. For INDEL, Clair3-Trio showed consistently better results than Clair3 and PEPPER. For INDEL, Clair3-Trio achieved an F1-score of 78.07% and 77.40%at 60x HG002 (child) and HG003 (parent) data, respectively. This is much higher than 72.45%, 75.02% in Clair3 (*p*-value < 0.05, two-tailed t-test) and 66.94%, 68.91% in PEPPER (*p*-value < 0.001, two-tailed t-test). These results verify the effectiveness of the Clair3-Trio model.

Comparing performance gain among members of the family trio, we found that the performance gain in the child (HG002) was much more profound than that in the parents (HG003 and HG004). For INDEL, Clair3-Trio achieved a +5.62% increment in the F1-score in the child compared to Clair3 at 60x, while the improvement dropped to +2.38% in the parents (**Supplementary Table 1**). The rationale is that for calling variants, the family trio provided more information about the child, which shares two haplotypes with parents, while each parent shares only one haplotype with the child.

### Assessing variant-calling accuracy in a family trio

Comprehensively evaluating variants across all family members using metrics like the number of Mendelian violations is important when calling variants in a family trio. In Mendelian inheritance violations, Clair3-Trio showed an order of magnitude fewer violations than Clair3 and PEPPER at 10x to 30x coverage. As shown in **Figure 3** and **Supplementary Table 1**, at 10x coverage, there were 7,072 Mendelian violations called from Clair3-Trio, while the number was 48,345 and 131,509 in Clair3 and PEPPER, respectively. At 60x, in contrast, there arewere 8,429 Mendelian violations called from Clair3-Trio, and 30,725 and 20,559 in Clair3 and PEPPER, respectively. Regarding the number of Mendelian violations found in Clair3-Trio at 60x, 70.3% and 29.7% of the calls were SNP and INDEL, respectively, indicating a large proportion of SNP and INDEL Mendelian violations recognised by Clair3-Trio. In contrast, for the number of *de novo* variants, Clair3-Trio has fewer false-positive (FP) *de novo* variants than other tools had. Clair3-Trio had 197 FPs at 60x data compared to 458 and 455 in Clair3 and PEPPER, respectively. However, Clair3-Trio found slightly fewer truepositive (TP) *de novo* variants, 33, compared to 35 in both Clair3 and PEPPER. We gathered all false-negative *de novo* variant cases of Clair3-Trio in **Supplementary Table 2** and present their alignment visualization in **Supplementary Figure 3**. More discussion about the Mendelian violations and *de novo* variant calling for Clair3-Trio are presented in the **Discussion** section.

**Figure 3.**
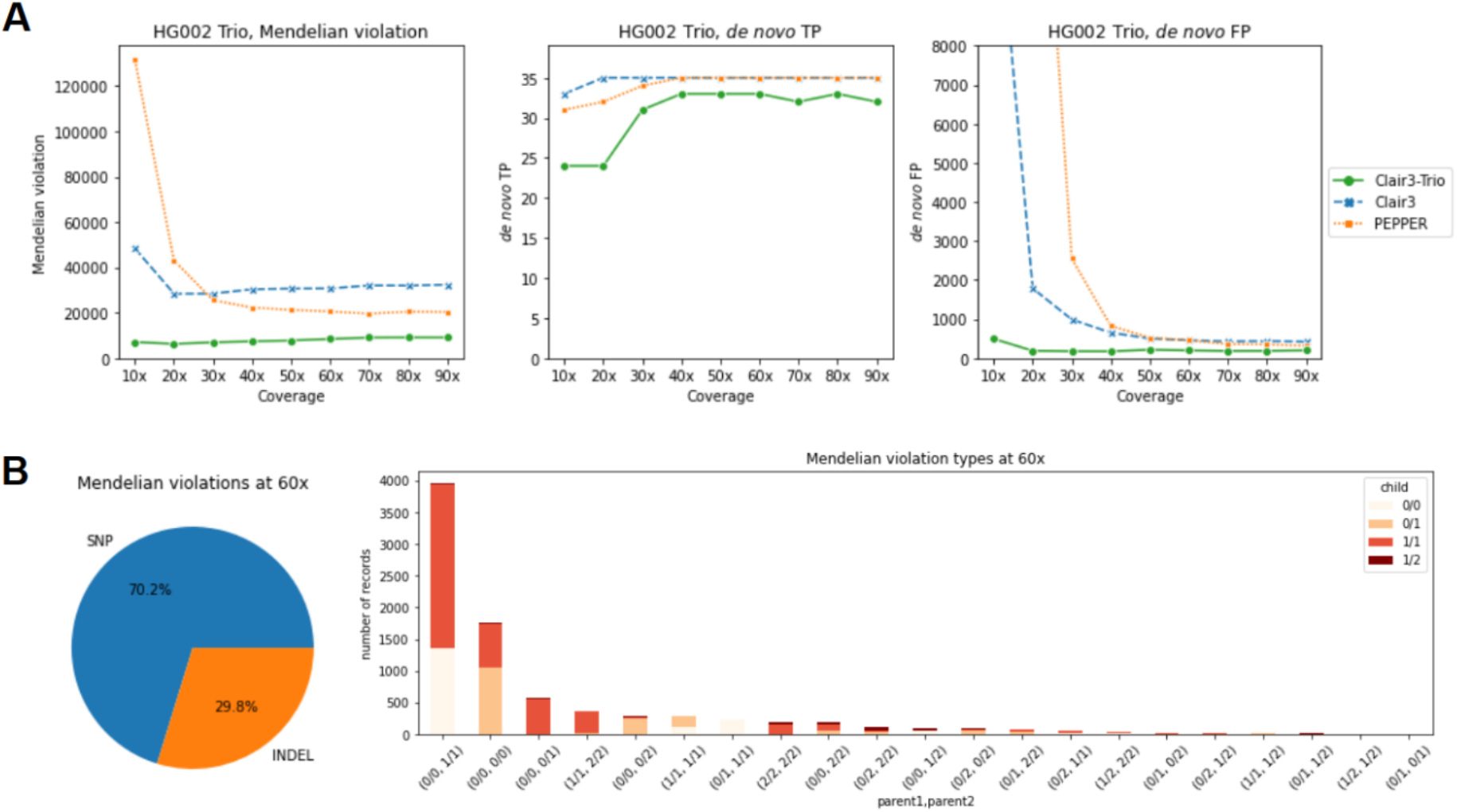
Trio benchmarking results on the GIAB trio. **A.** The number of Mendelian inheritance violations, *de novo* true positives and *de novo* false positives found in different tools. **B.** The breakdown of Mendelian violation rate of 60x from Clair3-Trio.

### Assessing the effect of varying parental coverage on variant-calling accuracy

When calling variants from trios, there are cases in which the data from the parents is only half or less of the coverage of the children. To assess the effect of low parental coverage on variant calling, we set the child sample to coverage of 60x and downsampled the sequencing data of parents from 60x into test ranges of 10x, 20x, 30x, 40x, 50x, and 60x. The test results are shown in **Figure 4** (SNP+INDEL), and further details are provided in **Supplementary Figure 4** (SNP), **Supplementary Figure 5** (INDEL), and **Supplementary Table 3**. For the child sample, the performance of Clair3-Trio is similar to that of Clair3 when the parent has very low coverage (10x) overall, indicating that 20x or more for parents is required for trio calling to improve the variant calling for the child. When the parents have half the child’s coverage (child 60x, parents 30x), Clair3-Trio achieved an overall F1-score of 96.92%, compared to 96.50% and 95.98% for Clair3 and PEPPER, respectively. Separating the results from SNP and INDEL, we found that Clair3-Trio outperformed the other tools when parents had coverage higher than 10x for SNP calling and coverage higher than 30x for INDEL calling. Further, in Clair3-Trio, there was a large improvement in the performance of low-coverage parent data when higher coverage for the child was provided (**Figure 4**). Clair3-Trio achieved a +6.02% increment in the F1-score in HG003 (10x parent sample) compared to Clair3. Furthermore, when parents had half the child’s coverage (60x for child and 30x for parents), Clair3-Trio had an F1-score of 96.54% for HG003, which is also higher than 95.83% in Clair3 and 95.07% in PEPPER. The improvement of Clair3-Trio on the trio data makes it useful for population genome projects in which better variant calling performance is expected for both parents and children.

**Figure 4.**
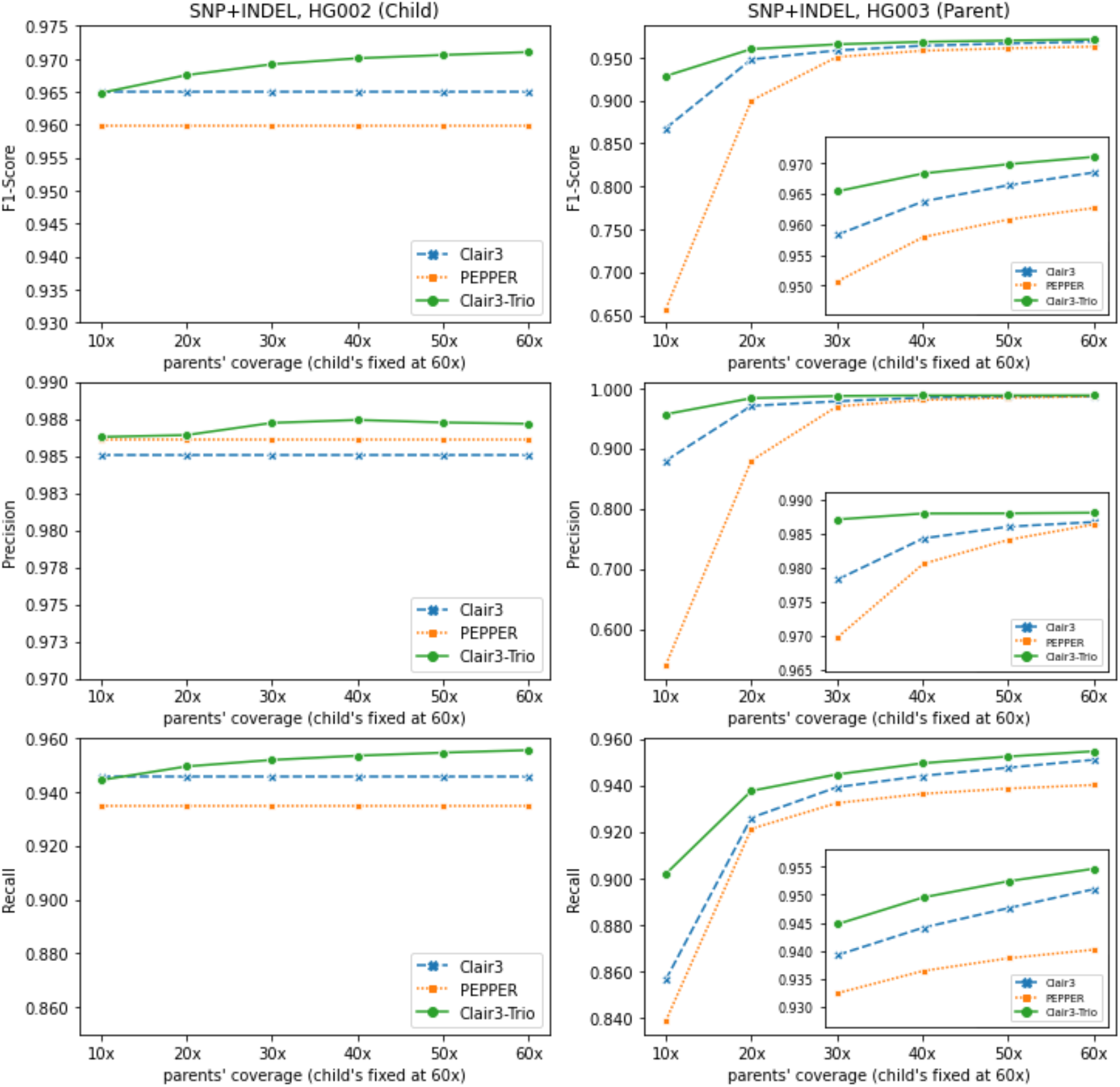
Overall benchmarking results on the GIAB trio when only the parent samples have diverse coverage. The SNP+INDEL’s F1-score, Precision and Recall of different tools at coverage from 10x to 60x on parent samples with the child’s coverage fixed at 60x.

### Building the Clair3-Trio model

#### Comparison of different architectures and model shape

We first categorized different methods based on their input and output information to generalize different methods for variant calling from family child-mother-father trio data. The One-to-One model inputs single sample information and outputs single sample variants. Clair3, PEPPER and Medaka are typical One-to-One models. The Trio-to-One model inputs data from three samples into the model and outputs single sample variants. For example, DeepTrio, which works with Illumina and PacBio HiFi data, is a typical Trio-to-One model. Finally, the Trio-to-Trio model inputs data from three samples into the model and outputs the three samples’ variants simultaneously. In Clair3-Trio, we built the first Trio-to-Trio model.

To compare the performance of different architectures, we ablated the input and output tensors of Clair3-Trio models accordingly to test as three architectures: One-to-One, Trio-to-One, and Trio-to-Trio models. The One-to-One model has single sample input and predicts single sample variants, as in Clair3 and PEPPER. The Trio-to-One model has information of three samples in its input, but predicts single sample variants in its model, as in DeepTrio. The Trio-to-Trio model is a native version of Clair3-Trio, which has three samples input and three samples output, but with deactivated MCVLoss and fine-tuning. We trained a single model for all architectures on chromosome 1 64x data from the GIAB HG002 trio and tested the performance on chromosome 20. For the Trio-to-One model, which is sample order specific, we trained two models separately to make predictions: a child model and a parent model. The benchmark results for the child as well as the number of Mendelian violations are in **Figure 5**.

**Figure 5.**
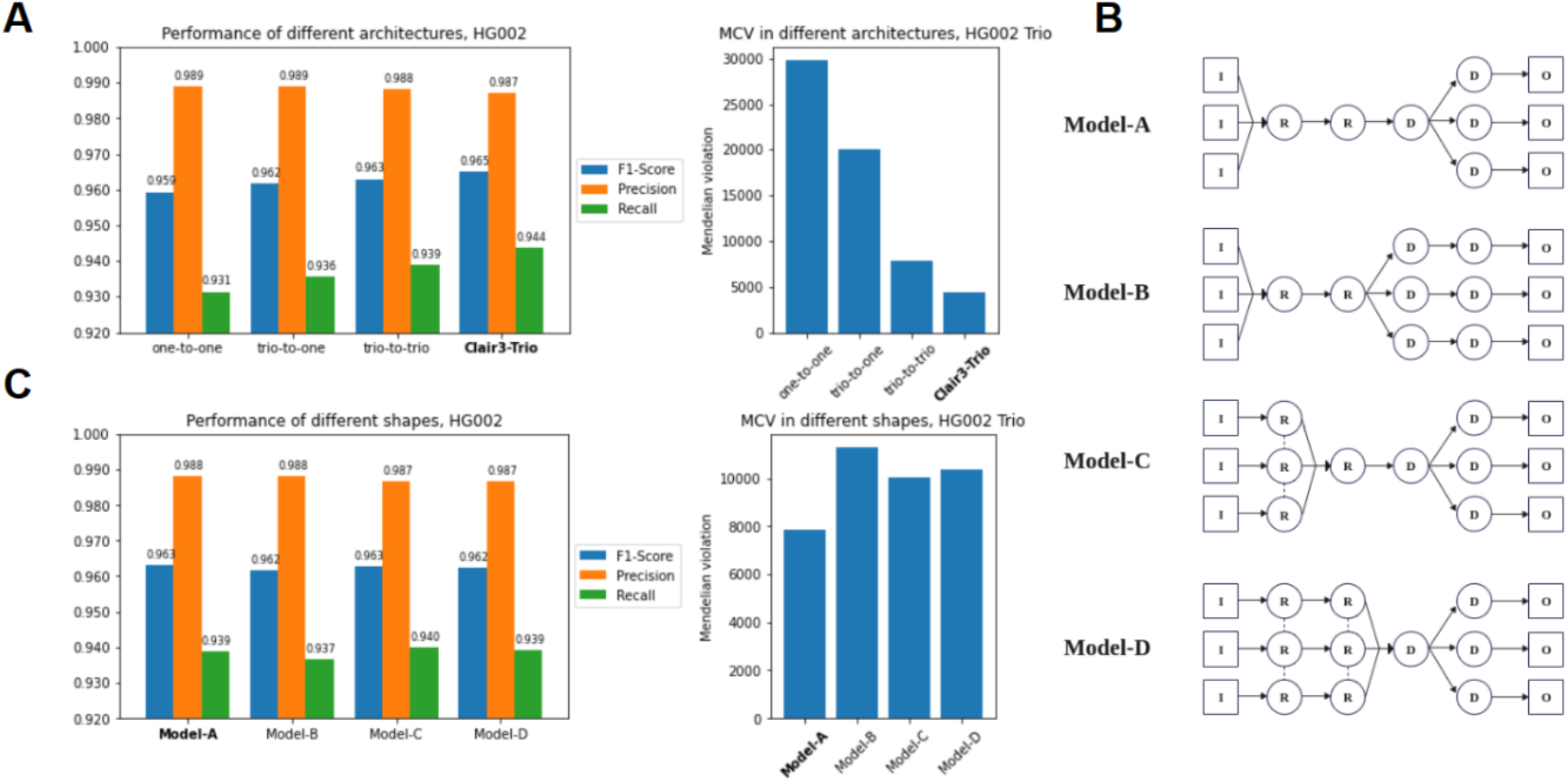
Comparison of different architectures and shapes for calling variants from trios. All results were trained on chr1 and tested on chr20. **A.** The performance of different model architecture, including one-to-one (Model-A), trio-to-one (Model-B), trio-to-trio (Model-C) and Clair3-Trio (Model-D, fine-tuned with MCVLoss) architecture; **B.** The schema of different trio-to-trio model shapes; and **C.** Comparison of different model shapes.

We found that including trios information in the model efficiently improves the variant calling performance overall, especially in terms of Mendelian inheritance violations (**Figure 5)**. Switching from One-to-One to Trio-to-One alone can boost the F1-score in the child by about +0.23%. The performance increment is consistent with the DeepTrio results [15]. The performance-boosting increased to +0.37% and +1.2% when the architecture was switched to Trio-to-Trio and Clair3-Trio (with MCVLoss and fine-tuning), respectively. For the child sample, the F1-score for Trio-to-Trio, One-to-One and Trio-to-One was 96.30%, 95.93%, and 96.16%, respectively. However, for the parent samples, Trio-to-Trio was only slightly better than One-to-One and Trio-to-One. We also found that the Trio-to-Trio architecture predicted many fewer Mendelian inheritance violation variants: 7,872 in the Trio-to-Trio model, 29,753 in the One-to-One model, and 20,016 in the Trio-to-One model.

To further explore the best architecture for the Trio-to-Trio model, we also evaluated the effect of using different model shapes. With three inputs and three outputs available, we developed multiple candidates for model shape, as illustrated in **Figure 5**: (1) Model-A, which inputs the information of all samples into Resblock and divides the last dense layer to give three outputs; (2) Model-B, which inputs the information of all samples into Resblock divided at all dense layers; (3) Model-C, which inputs single sample information into shared Resblock, and divides the last dense layer to generate three outputs; and (4) Model-D, which shares multiple Resblock from a single input and divides the last dense layer to generate three outputs. We found that Model-A and Model-C achieved a similar F1-score (96.30% for Model-A and 96.26% for Model-C) in the child sample to that in Model-B (96.18%) and Model-D (96.25%), but Model-A had many fewer Mendelian violation predictions than the other models (7,872 compared to 11,278, 10,053 and 10,370, respectively, in the other shapes). For this reason, we selected Model-A as the best shape for the Trio-to-Trio architecture.

#### Finetuning with MCVLoss

The **MCVLoss** (**M**endelian Inheritance **C**onstraint **V**iolation **Loss**) function is designed to improve variant calling in trios by leveraging the explicit encoding of the priors of the Mendelian inheritance in trios. We found that MCVLoss can effectively reduce Mendelian violation prediction in variant calling. However, the prediction is better accompanied with fine-tuning techniques, in which we train a Clair3-Trio model in two steps: (1) training Clair3-Trio without MCVLoss with the default learning rate (1e-3 in our setting), and (2) fine-tuning the trained Clair3-Trio model with MCVLoss with a lower learning rate (1e-5 in our setting). When using the fine-tuning technique alone, the F1-score from HG002, HG003 and HG004 had a performance boost of +0.2% (**Table 1**). We got the best results when combining finetuning and MCVLoss with the +0.2% F1-score increment and a Mendelian violations reduction from 7,872 to 4,352.

**Table 1.** Benchmarking results from fine-tuning with MCVLoss. All results were tested at 64x data. FT: fine-tune; “# of MCV”: number of Mendelian inheritance violations in predicted variants.

| FT | MCV-Loss | Sample | Overall |  |  | SNP |  |  | INDEL |  |  | # of MCV |
| --- | --- | --- | --- | --- | --- | --- | --- | --- | --- | --- | --- | --- |
|  |  |  | Precision | Recall | F1-Score | Precision | Recall | F1-Score | Precision | Recall | F1-Score |  |
| N | N | HG002 | 98.83% | 93.90% | 96.30% | 99.72% | 99.57% | 99.64% | 90.02% | 57.92% | 70.49% | 7872 |
|  |  | HG003 | 98.72% | 94.12% | 96.36% | 99.65% | 99.54% | 99.59% | 89.34% | 58.35% | 70.59% |  |
|  |  | HG004 | 98.84% | 93.93% | 96.32% | 99.70% | 99.58% | 99.64% | 90.01% | 57.06% | 69.84% |  |
| Y | N | HG002 | 98.79% | 94.31% | 96.50% | 99.67% | 99.68% | 99.67% | 90.45% | 60.21% | 72.29% | 4885 |
|  |  | HG003 | 98.83% | 94.42% | 96.58% | 99.61% | 99.64% | 99.63% | 90.99% | 59.96% | 72.29% |  |
|  |  | HG004 | 98.91% | 94.22% | 96.51% | 99.69% | 99.66% | 99.67% | 91.06% | 58.78% | 71.44% |  |
| N | Y | HG002 | 99.02% | 93.59% | 96.23% | 99.72% | 99.56% | 99.64% | 91.69% | 55.69% | 69.30% | 4754 |
|  |  | HG003 | 98.87% | 93.95% | 96.34% | 99.64% | 99.61% | 99.63% | 90.64% | 56.58% | 69.67% |  |
|  |  | HG004 | 98.83% | 93.95% | 96.32% | 99.72% | 99.61% | 99.66% | 89.64% | 57.05% | 69.72% |  |
| Y | Y | HG002 | 98.72% | 94.37% | 96.50% | 99.67% | 99.68% | 99.68% | 89.75% | 60.70% | 72.42% | 4352 |
|  |  | HG003 | 98.77% | 94.46% | 96.57% | 99.61% | 99.63% | 99.62% | 90.47% | 60.31% | 72.37% |  |
|  |  | HG004 | 98.88% | 94.27% | 96.52% | 99.69% | 99.66% | 99.67% | 90.83% | 59.14% | 71.64% |  |

We also evaluated the effect of using a different *α* rate in MCVLoss (**Table 2**). The *α* rate in MCVLoss controls the weighting in terms of loss function, as in formula (9). We observed that increasing the *α* rate efficiently decreases the number of Mendelian inheritance violations, but slightly decreases the overall performance based on the F1-score. We found the *α* rate of 1 to be the best setting for MCVLoss, which balances the F1-score and the number of Mendelian inheritance violations metrics.

**Table 2.** Benchmarking results of different *α* rate in MCVLoss. All results were tested at 64x data. “# of MCV”: number of Mendelian inheritance violations in predicted variants.

| $\alpha$ | Sample | Overall | | | SNP | | | INDEL | | | # of MCV |
| --- | --- | --- | --- | --- | --- | --- | --- | --- | --- | --- | --- |
|  |  | Precision | Recall | F1-Score | Precision | Recall | F1-Score | Precision | Recall | F1-Score |  |
| 0 | HG002 | 98.79% | 94.31% | 96.50% | 99.67% | 99.68% | 99.67% | 90.45% | 60.21% | 72.29% | 4885 |
|  | HG003 | 98.83% | 94.42% | 96.58% | 99.61% | 99.64% | 99.63% | 90.99% | 59.96% | 72.29% |  |
|  | HG004 | 98.91% | 94.22% | 96.51% | 99.69% | 99.66% | 99.67% | 91.06% | 58.78% | 71.44% |  |
| 0.1 | HG002 | 98.77% | 94.33% | 96.50% | 99.67% | 99.68% | 99.68% | 90.19% | 60.38% | 72.33% | 4990 |
|  | HG003 | 98.83% | 94.42% | 96.57% | 99.62% | 99.63% | 99.63% | 90.91% | 59.97% | 72.27% |  |
|  | HG004 | 99.12% | 94.06% | 96.52% | 99.71% | 99.65% | 99.68% | 92.93% | 57.61% | 71.13% |  |
| 1 | HG002 | 98.72% | 94.37% | 96.50% | 99.67% | 99.68% | 99.68% | 89.75% | 60.70% | 72.42% | 4352 |
|  | HG003 | 98.77% | 94.46% | 96.57% | 99.61% | 99.63% | 99.62% | 90.47% | 60.31% | 72.37% |  |
|  | HG004 | 98.88% | 94.27% | 96.52% | 99.69% | 99.66% | 99.67% | 90.83% | 59.14% | 71.64% |  |
| 10 | HG002 | 98.87% | 94.17% | 96.46% | 99.69% | 99.64% | 99.66% | 90.90% | 59.47% | 71.90% | 3926 |
|  | HG003 | 98.63% | 94.49% | 96.51% | 99.59% | 99.63% | 99.61% | 89.30% | 60.53% | 72.15% |  |
|  | HG004 | 98.96% | 94.19% | 96.51% | 99.68% | 99.66% | 99.67% | 91.66% | 58.47% | 71.39% |  |

## Discussion

In this paper, we introduced Clair3-Trio, a high-performance Nanopore long-read variant caller in family trios with a Trio-to-Trio deep neural network. Clair3-Trio is the first family trio variant caller tailored for Nanopore long-read data with a Trio-to-Trio deep neural network model and MCVLoss. In our experiments, Clair3-Trio significantly outperformed current state-of-the-art methods on trio variant calling in terms of F1-score and the number of Mendelian inheritance violations in all three samples from a trio. We also demonstrated that the architecture of the Trio-to-Trio model is much more accurate than the One-to-One and Trio-to-One model. The source code and the results of this study are publicly available on GitHub.

We found that most of the Mendelian violationcases from Clair3-Trio (68.6%) for parent-1, parent-2 and child, respectively, are: (0/0, 1/1, 0/0), (0/0, 1/1, 1/1), (0/0, 0/0, 0/1), (0/0, 0/0, 1/1) (**Figure 3B**). All these violations are prone to be found when there is a switch between heterozygosity and homozygosity in a single trio sample at a site. For example, in the case of Mendelian violations (0/0, 1/1, 0/0), the switch between heterozygous and homozygous in any member’s calling changes the variant calling to a non-Mendelian inheritance violation call. As all members have a chance of being miscalled, these cases remain a challenge even when trio data is available.

Clair3-Trio has high performance overall, but it has fewer *de novo* variants predicted than Clair3 and PEPPER. The drop in TP of *de novo* variants is expected, as Clair3-Trio is designed to predict variants by leveraging information from family trios that favor having fewer Mendelian violations in their prediction. For detecting *de novo* variants that do not follow Mendelian inheritance, One-to-One model-based methods such as Clair3 and PEPPER can be used to supplement Clair3-Trio.

There are some challenges and future works needed regarding trio variant calling from ONT data. Experiments show that Clair3-Trio’s improvement over state-of-the-art methods is profound when the trio data has similar coverage among family members, but it only marginally improves with calling variants from different data coverage (such as child coverage of 60x and parent coverage of 10x). These results leave room for further improvement in trio calling in diverse coverage applications. The current model is trained with multiple coverage down-sampled from the full coverage, but only with the coverage of the child kept equal to or larger than that of the parents, and not the cartesian product of the down-sampled coverage of the three samples. This is a practical decision to reduce the amount of training data and since the coverage of the child in a trio is usually higher than that of the parents. However, this may also challenge Clair3-Trio when the coverage of parents exceeds that of the child. An improved training scheme is expected to handle the large amount of training data when all coverage combinations are used. On the other hand, there is a research gap in applying variant calling in the human sex chromosome region. The current training and testing was constrained to the autosome region, which assumes that the variants are diplotypes and inherited from one of the parents. However, on the sex chromosome, the assumption is unheld when calling variants in the child’s Y chromosome, which is a haplotype and is obtained only from the father’s side. Currently, there are no tools available for calling variants in the sex chromosome region with the family information from ONT data. We need a new design for calling variants from the sex chromosome region to fill this research gap. In the future, we would like to design a heuristic approach to solve the question: if the child is female, use Clair3-Trio directly at the sex chromosome; if the child is male, use Clair3-Trio to call variants in the pseudoautosomal regions (PAR1 and PAR2) of the sex chromosome and build a tailored haplotype model to call variants in the remaining regions.

## Data availability

Clair3-Trio is open-source software (BSD 3-Clause license), hosted by GitHub at https://github.com/HKU-BAL/Clair3-Trio. The 1) links to the reference genomes, true variants, benchmarking materials, and ONT data, and 2) commands and parameters used in this study are available in the **Supplementary Notes**. All analysis outputs, including the VCFs and running logs, are available at http://www.bio8.cs.hku.hk/clair3_trio/analysis_result.

## Acknowledgments

R.L. was supported by the GRF [17113721] of the HKSAR Government, General Program [JCYJ20210324134405015] of the Shenzhen Municipal Government, and URC fund at HKU.

## Author contributions

R. L. conceived the study. J. S. and R. L. designed the algorithms, implemented Clair3-Trio, and wrote the paper. All authors evaluated the results and revised the manuscript.

## Key points

Developed a Trio-to-Trio model to predict trio variants in ONT data.

Introduced a novel loss function, MCVLoss, to model Mendelian inheritance in trio data.

Demonstrated that the Clair3-Trio model trained on GIAB data improves variant calling in trio data.

Demonstrated that Trio-to-Trio models can efficiently decrease Mendelian inheritance violations compared to One-to-One and Trio-to-One models.

