## Supplementary Materials for "Clair3-Trio: high-performance Nanopore long-read variant calling in family trios with Trio-to-Trio deep neural networks"

### Supplementary Figures

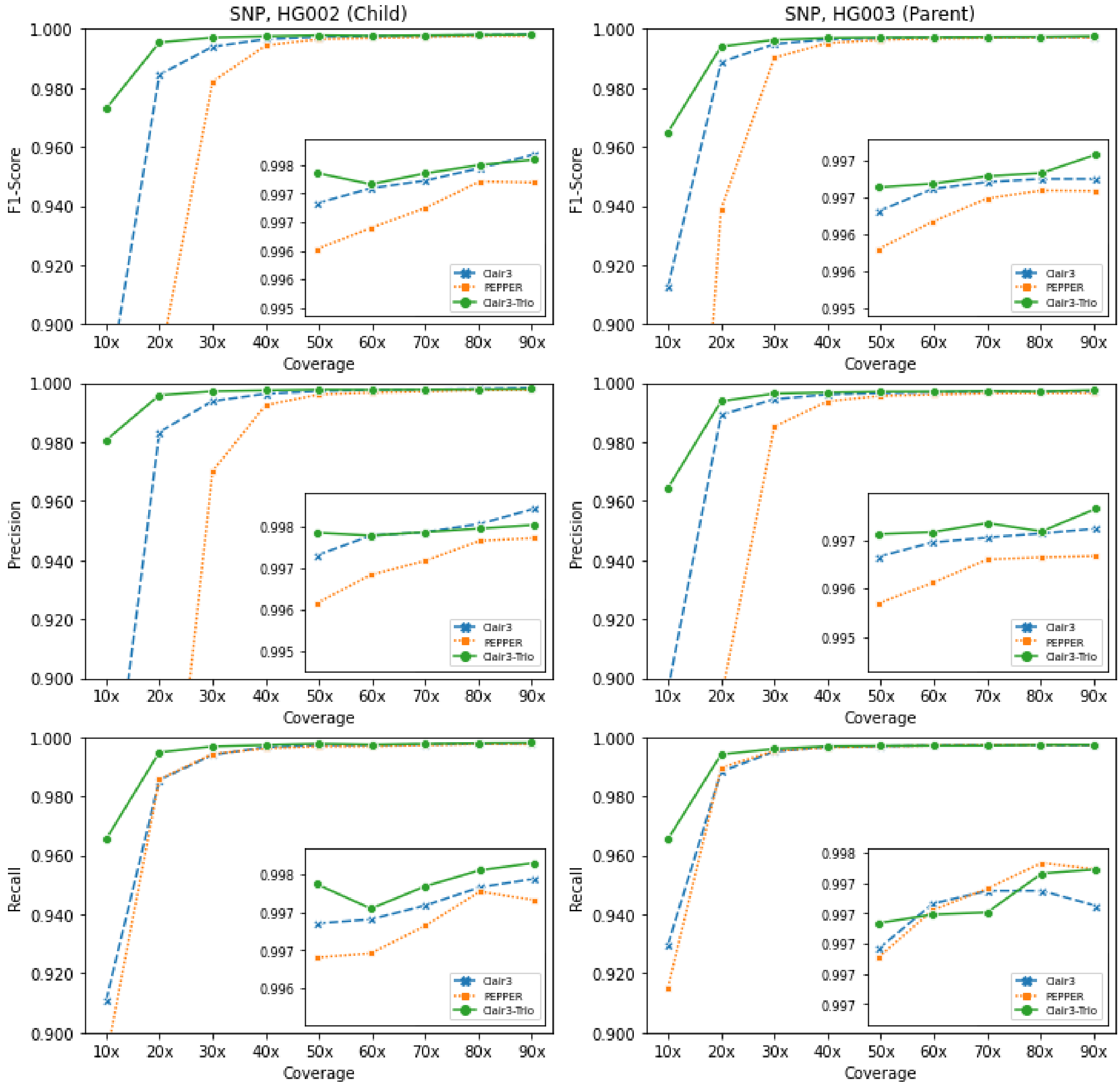

Supplementary Figure 1. SNP benchmarking results on the GIAB trio.

The SNP's F1-score, Precision and Recall for HG002 (child, left) and HG003 (parent, right) of different tools at coverages from 10x to 90x.

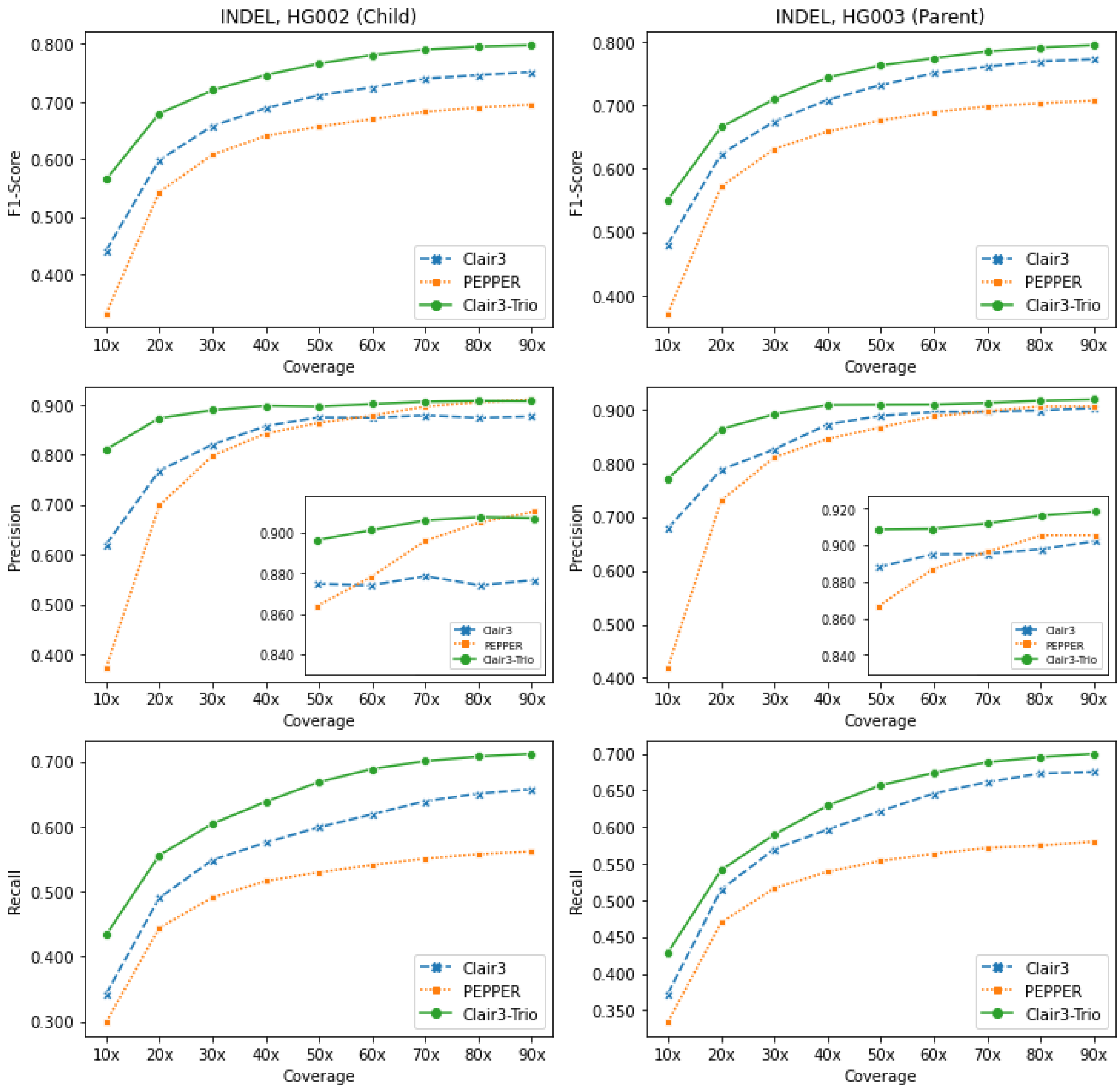

### Supplementary Figure 2. INDEL benchmarking results on the GIAB trio.

The INDEL's F1-score, Precision and Recall for HG002 (child, left) and HG003 (parent, right) of different tools at coverages from 10x to 90x.

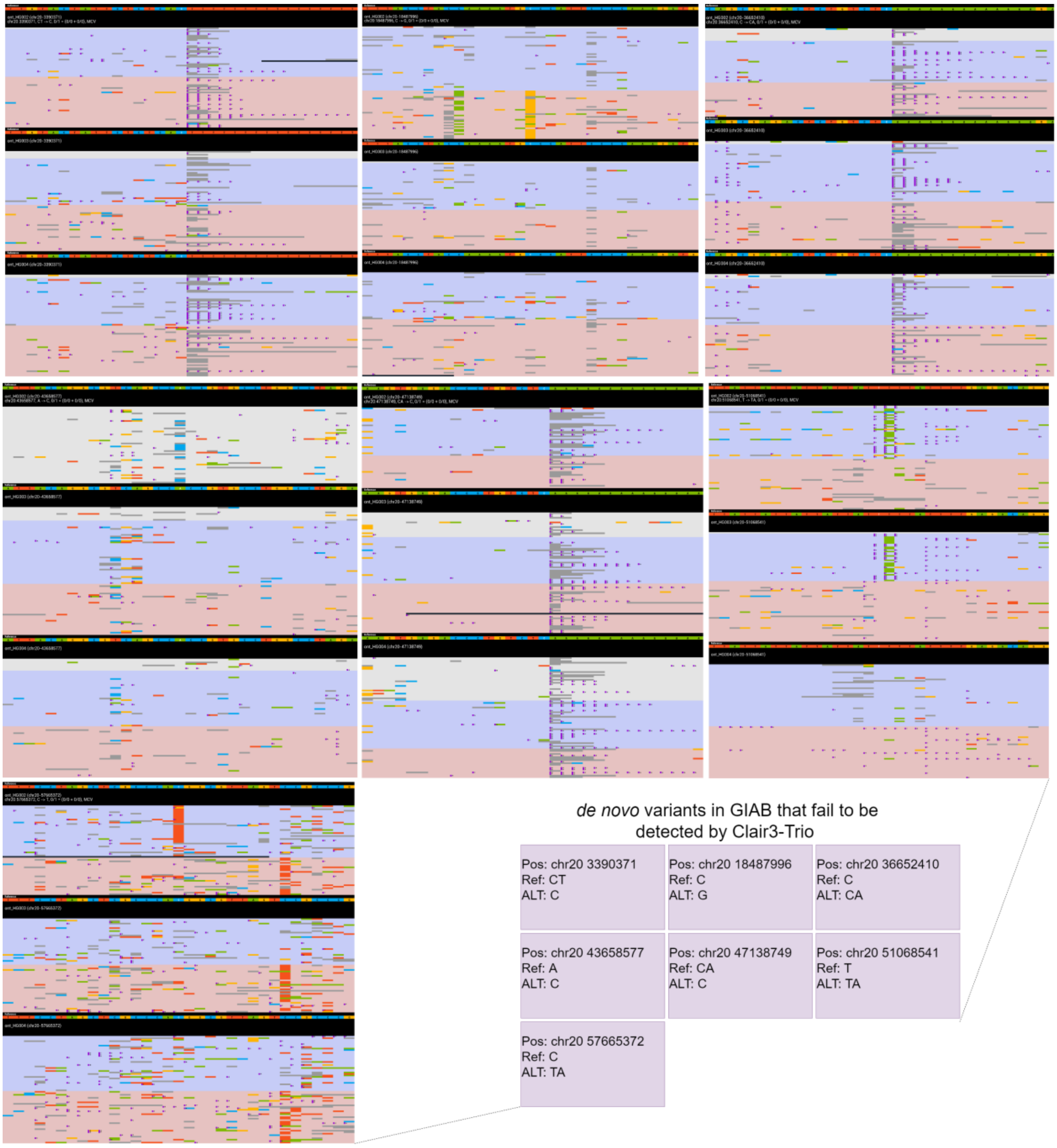

Supplementary Figure 3. Phased alignment visualization for *de novo* variants that fail to be detected by Clair3-Trio.

For each visualization, from top to bottom is alignment for HG002 (child), HG003 (parent1), HG004 (parent2). The A, C, G, T, insertion, and deletion are in green, blue, yellow, red, purple dots, and grey, respectively.

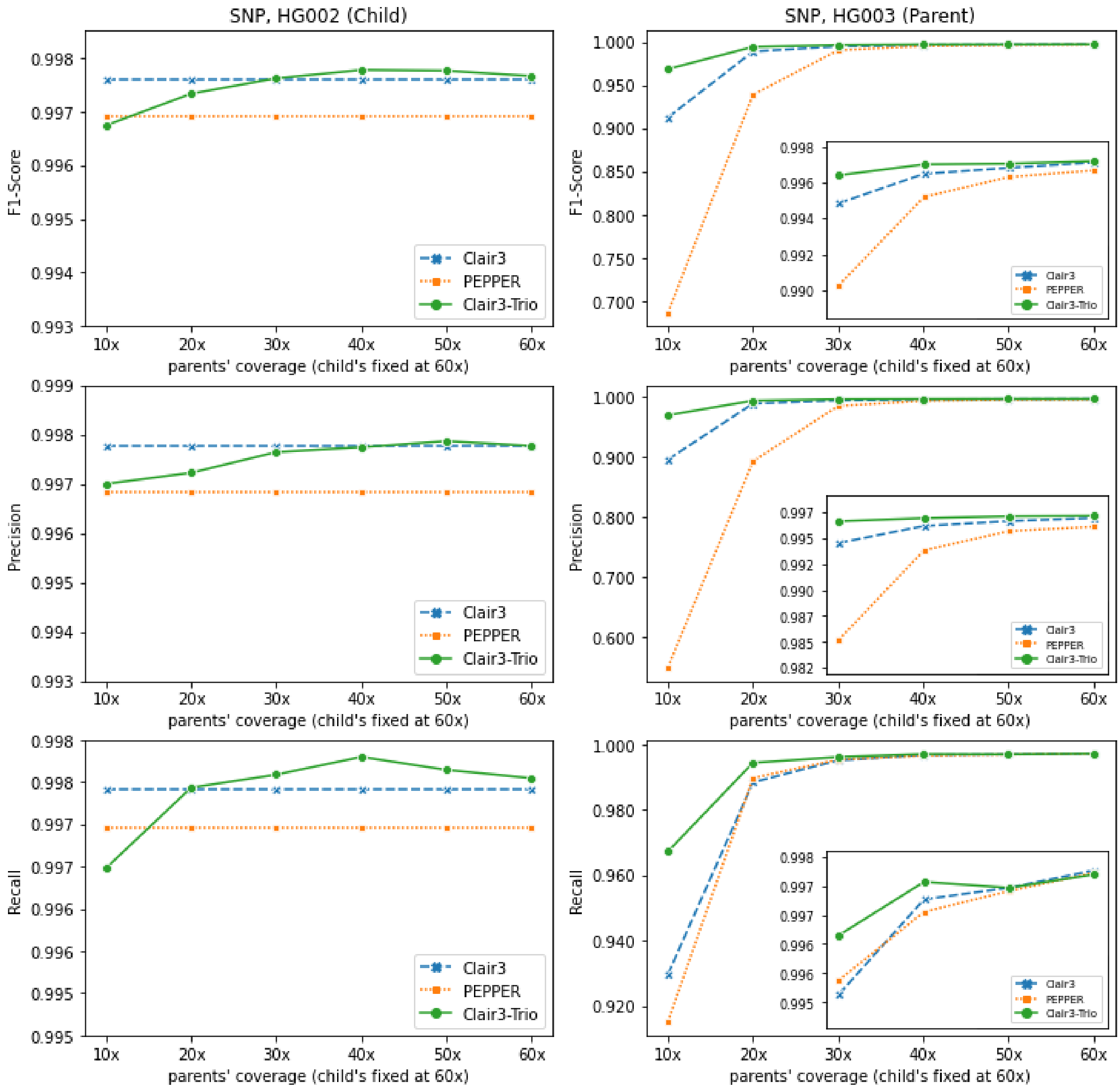

Supplementary Figure 4. SNP benchmarking results on the GIAB trio when only parents have diverse depth.

The SNP's F1-score, Precision and Recall for HG003 (parent) of different tools at coverages from 10x to 60x while child's coverage fixed at 60x is shown.

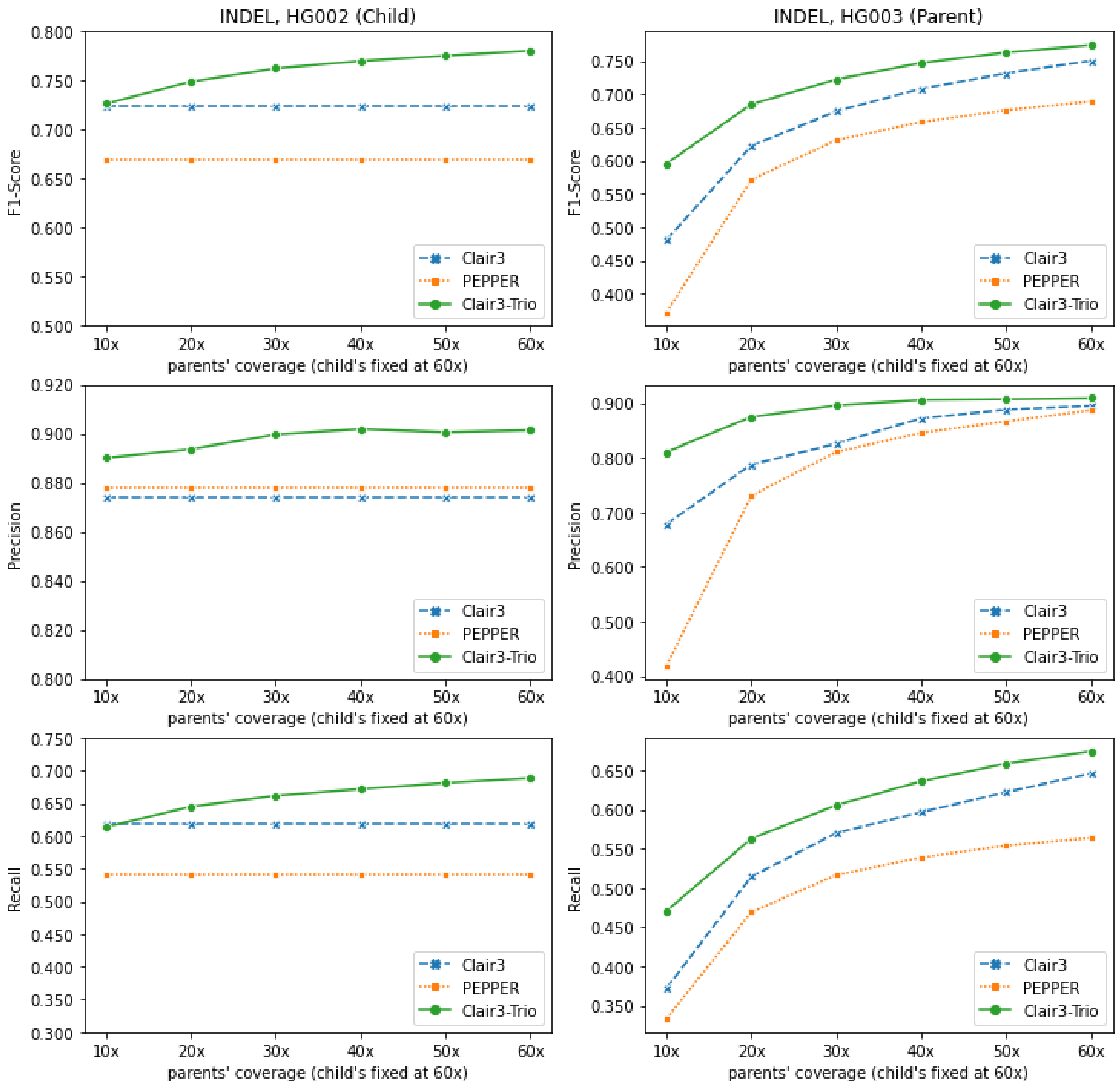

Supplementary Figure 5. INDEL benchmarking results on the GIAB trio when only parents have diverse depth.

The INDEL's F1-score, Precision and Recall for HG003 (parent) of different tools at coverages from 10x to 60x while child's coverage fixed at 60x is shown.

### Supplementary Tables

#### Supplementary Table 1. HG002 trio multiple-coverage benchmarking results.

| | | | Overall | | | SNP | | | INDEL | | | Insertion | | | Deletion | | | | | |
|---|---|---|---|---|---|---|---|---|---|---|---|---|---|---|---|---|---|---|---|---|
| Coverage | Tool | Sample | Precision | Recall | F1-Score | Precision | Recall | F1-Score | Precision | Recall | F1-Score | Precision | Recall | F1-Score | Precision | Recall | F1-Score | # of MCV | *de novo* TP | *de novo* FP |
| 10x | Clair3 | HG002 | 82.24% | 83.33% | 82.78% | 83.89% | 91.08% | 87.33% | 62.00% | 34.25% | 44.12% | 66.85% | 34.89% | 45.85% | 57.95% | 33.65% | 42.58% | 48345 | 33 | 12979 |
| | Clair3 | HG003 | 87.95% | 85.63% | 86.77% | 89.59% | 92.96% | 91.24% | 67.83% | 37.24% | 48.08% | 70.41% | 39.19% | 50.35% | 65.34% | 35.42% | 45.94% | | | |
| | Clair3 | HG004 | 86.98% | 84.91% | 85.93% | 88.70% | 92.40% | 90.51% | 66.02% | 36.14% | 46.71% | 66.12% | 37.47% | 47.83% | 65.92% | 34.92% | 45.66% | | | |
| 20x | Clair3 | HG002 | 96.34% | 91.80% | 94.02% | 98.34% | 98.55% | 98.45% | 76.76% | 49.01% | 59.82% | 77.85% | 49.76% | 60.71% | 75.73% | 48.31% | 58.99% | 28411 | 35 | 1787 |
| | Clair3 | HG003 | 97.07% | 92.61% | 94.79% | 98.92% | 98.84% | 98.88% | 78.76% | 51.46% | 62.25% | 78.48% | 53.16% | 63.38% | 79.04% | 49.88% | 61.16% | | | |
| | Clair3 | HG004 | 97.07% | 92.56% | 94.76% | 99.11% | 98.92% | 99.02% | 77.39% | 51.07% | 61.53% | 75.49% | 52.71% | 62.08% | 79.35% | 49.56% | 61.01% | | | |
| 30x | Clair3 | HG002 | 97.70% | 93.33% | 95.47% | 99.39% | 99.40% | 99.40% | 82.05% | 54.86% | 65.75% | 83.51% | 55.78% | 66.89% | 80.68% | 54.01% | 64.70% | 28434 | 35 | 990 |
| | Clair3 | HG003 | 97.82% | 93.92% | 95.83% | 99.45% | 99.51% | 99.48% | 82.57% | 57.00% | 67.44% | 82.08% | 58.75% | 68.48% | 83.07% | 55.36% | 66.44% | | | |
| | Clair3 | HG004 | 98.02% | 93.80% | 95.86% | 99.66% | 99.58% | 99.62% | 82.69% | 56.14% | 66.88% | 80.50% | 58.18% | 67.55% | 84.97% | 54.26% | 66.23% | | | |
| 40x | Clair3 | HG002 | 98.28% | 93.92% | 96.05% | 99.64% | 99.67% | 99.65% | 85.73% | 57.51% | 68.84% | 86.75% | 58.44% | 69.83% | 84.78% | 56.64% | 67.91% | 30352 | 35 | 652 |
| | Clair3 | HG003 | 98.43% | 94.41% | 96.38% | 99.62% | 99.68% | 99.65% | 87.21% | 59.65% | 70.84% | 87.49% | 61.69% | 72.36% | 86.94% | 57.74% | 69.39% | | | |
| | Clair3 | HG004 | 98.55% | 94.27% | 96.37% | 99.76% | 99.69% | 99.72% | 87.27% | 58.95% | 70.37% | 85.18% | 60.95% | 71.06% | 89.43% | 57.11% | 69.70% | | | |
| 50x | Clair3 | HG002 | 98.51% | 94.31% | 96.36% | 99.73% | 99.74% | 99.73% | 87.48% | 59.89% | 71.10% | 88.04% | 60.07% | 71.41% | 86.97% | 59.72% | 70.81% | 30674 | 35 | 502 |
| | Clair3 | HG003 | 98.60% | 94.76% | 96.64% | 99.67% | 99.70% | 99.68% | 88.81% | 62.19% | 73.15% | 89.03% | 64.38% | 74.73% | 88.59% | 60.14% | 71.64% | | | |
| | Clair3 | HG004 | 98.74% | 94.70% | 96.68% | 99.80% | 99.74% | 99.77% | 89.04% | 61.81% | 72.97% | 87.23% | 63.88% | 73.75% | 90.90% | 59.92% | 72.22% | | | |
| 60x | Clair3 | HG002 | 98.51% | 94.58% | 96.50% | 99.78% | 99.74% | 99.76% | 87.41% | 61.86% | 72.45% | 88.53% | 61.86% | 72.83% | 86.39% | 61.86% | 72.10% | 30725 | 35 | 458 |
| | Clair3 | HG003 | 98.67% | 95.10% | 96.85% | 99.70% | 99.73% | 99.71% | 89.51% | 64.57% | 75.02% | 90.06% | 66.06% | 76.21% | 88.96% | 63.18% | 73.89% | | | |
| | Clair3 | HG004 | 98.88% | 95.08% | 96.94% | 99.81% | 99.76% | 99.79% | 90.60% | 64.56% | 75.39% | 89.38% | 66.95% | 76.56% | 91.84% | 62.36% | 74.28% | | | |
| 70x | Clair3 | HG002 | 98.53% | 94.87% | 96.67% | 99.79% | 99.76% | 99.77% | 87.86% | 63.88% | 73.97% | 89.66% | 63.59% | 74.41% | 86.27% | 64.14% | 73.58% | 32041 | 35 | 428 |
| | Clair3 | HG003 | 98.66% | 95.32% | 96.96% | 99.71% | 99.74% | 99.72% | 89.54% | 66.13% | 76.08% | 90.62% | 67.23% | 77.19% | 88.52% | 65.11% | 75.03% | | | |
| | Clair3 | HG004 | 98.94% | 95.29% | 97.08% | 99.84% | 99.78% | 99.81% | 91.08% | 66.08% | 76.59% | 90.04% | 68.34% | 77.70% | 92.12% | 64.02% | 75.54% | | | |
| 80x | Clair3 | HG002 | 98.47% | 95.05% | 96.73% | 99.81% | 99.78% | 99.79% | 87.41% | 65.04% | 74.58% | 90.02% | 64.55% | 75.19% | 85.14% | 65.49% | 74.04% | 31996 | 35 | 438 |
| | Clair3 | HG003 | 98.68% | 95.47% | 97.05% | 99.71% | 99.74% | 99.72% | 89.79% | 67.31% | 76.94% | 91.30% | 67.93% | 77.90% | 88.39% | 66.73% | 76.05% | | | |
| | Clair3 | HG004 | 99.01% | 95.51% | 97.23% | 99.85% | 99.80% | 99.82% | 91.87% | 67.61% | 77.90% | 91.00% | 69.63% | 78.89% | 92.74% | 65.76% | 76.95% | | | |
| 90x | Clair3 | HG002 | 98.53% | 95.15% | 96.81% | 99.84% | 99.79% | 99.82% | 87.67% | 65.72% | 75.12% | 90.66% | 64.76% | 75.55% | 85.12% | 66.60% | 74.73% | 32368 | 35 | 420 |
| | Clair3 | HG003 | 98.74% | 95.49% | 97.08% | 99.72% | 99.73% | 99.72% | 90.24% | 67.51% | 77.24% | 91.82% | 68.39% | 78.39% | 88.77% | 66.69% | 76.16% | | | |
| | Clair3 | HG004 | 99.05% | 95.62% | 97.30% | 99.86% | 99.81% | 99.83% | 92.16% | 68.28% | 78.45% | 91.47% | 70.10% | 79.37% | 92.84% | 66.61% | 77.57% | | | |
| 10x | PEPPER | HG002 | 41.37% | 81.04% | 54.77% | 41.61% | 89.10% | 56.73% | 37.28% | 29.92% | 33.20% | 32.93% | 32.90% | 32.91% | 43.72% | 27.17% | 33.51% | 131509 | 31 | 87600 |
| | PEPPER | HG003 | 54.01% | 83.86% | 65.70% | 54.89% | 91.51% | 68.62% | 41.90% | 33.37% | 37.15% | 37.34% | 37.35% | 37.35% | 48.92% | 29.65% | 36.92% | | | |
| | PEPPER | HG004 | 50.55% | 83.40% | 62.95% | 51.21% | 91.22% | 65.60% | 40.95% | 32.50% | 36.24% | 35.56% | 36.33% | 35.94% | 49.50% | 28.97% | 36.55% | | | |

| | | | | | | | | | | | | | | | | | | | | |
|---|---|---|---|---|---|---|---|---|---|---|---|---|---|---|---|---|---|---|---|---|
| 20x | PEPPER | HG002 | 79.59% | 91.19% | 84.99% | 80.40% | 98.57% | 88.56% | 69.81% | 44.40% | 54.28% | 68.85% | 47.19% | 56.00% | 70.84% | 41.82% | 52.59% | 43023 | 32 | 17434 |
| | PEPPER | HG003 | 87.96% | 92.13% | 90.00% | 89.30% | 98.97% | 93.89% | 73.03% | 46.95% | 57.15% | 71.07% | 51.42% | 59.67% | 75.35% | 42.77% | 54.56% | | | |
| | PEPPER | HG004 | 87.86% | 91.92% | 89.84% | 89.28% | 99.00% | 93.89% | 71.98% | 45.77% | 55.96% | 67.84% | 49.75% | 57.41% | 77.07% | 42.11% | 54.46% | | | |
| 30x | PEPPER | HG002 | 95.50% | 92.58% | 94.02% | 97.02% | 99.44% | 98.21% | 79.80% | 49.08% | 60.77% | 78.99% | 52.03% | 62.74% | 80.65% | 46.34% | 58.86% | 25540 | 34 | 2551 |
| | PEPPER | HG003 | 96.97% | 93.24% | 95.07% | 98.52% | 99.54% | 99.03% | 81.08% | 51.65% | 63.10% | 78.89% | 55.90% | 65.44% | 83.63% | 47.68% | 60.73% | | | |
| | PEPPER | HG004 | 97.14% | 93.07% | 95.06% | 98.74% | 99.60% | 99.17% | 80.69% | 50.50% | 62.12% | 76.23% | 54.35% | 63.45% | 86.04% | 46.96% | 60.76% | | | |
| 40x | PEPPER | HG002 | 97.92% | 93.08% | 95.44% | 99.26% | 99.62% | 99.44% | 84.26% | 51.61% | 64.01% | 84.48% | 54.58% | 66.32% | 84.03% | 48.85% | 61.78% | 22279 | 35 | 826 |
| | PEPPER | HG003 | 98.05% | 93.64% | 95.79% | 99.38% | 99.66% | 99.52% | 84.53% | 53.90% | 65.83% | 82.31% | 58.18% | 68.17% | 87.08% | 49.90% | 63.45% | | | |
| | PEPPER | HG004 | 98.16% | 93.47% | 95.76% | 99.56% | 99.70% | 99.63% | 84.02% | 52.90% | 64.92% | 80.30% | 56.80% | 66.53% | 88.37% | 49.31% | 63.30% | | | |
| 50x | PEPPER | HG002 | 98.43% | 93.32% | 95.81% | 99.62% | 99.69% | 99.65% | 86.37% | 52.95% | 65.65% | 86.29% | 55.76% | 67.75% | 86.44% | 50.34% | 63.63% | 21267 | 35 | 524 |
| | PEPPER | HG003 | 98.41% | 93.86% | 96.08% | 99.57% | 99.69% | 99.63% | 86.66% | 55.40% | 67.59% | 84.83% | 59.53% | 69.96% | 88.71% | 51.54% | 65.20% | | | |
| | PEPPER | HG004 | 98.50% | 93.70% | 96.04% | 99.68% | 99.73% | 99.70% | 86.54% | 54.39% | 66.80% | 82.91% | 58.30% | 68.46% | 90.74% | 50.79% | 65.13% | | | |
| 60x | PEPPER | HG002 | 98.61% | 93.48% | 95.98% | 99.68% | 99.70% | 99.69% | 87.81% | 54.09% | 66.94% | 86.99% | 56.58% | 68.56% | 88.67% | 51.78% | 65.38% | 20559 | 35 | 455 |
| | PEPPER | HG003 | 98.64% | 94.02% | 96.27% | 99.61% | 99.72% | 99.67% | 88.70% | 56.35% | 68.91% | 86.80% | 60.31% | 71.17% | 90.82% | 52.65% | 66.66% | | | |
| | PEPPER | HG004 | 98.74% | 93.89% | 96.26% | 99.76% | 99.77% | 99.77% | 88.37% | 55.61% | 68.27% | 85.10% | 59.24% | 69.86% | 92.07% | 52.28% | 66.69% | | | |
| 70x | PEPPER | HG002 | 98.81% | 93.65% | 96.16% | 99.72% | 99.73% | 99.72% | 89.63% | 55.07% | 68.22% | 89.13% | 57.61% | 69.98% | 90.14% | 52.72% | 66.53% | 19583 | 35 | 359 |
| | PEPPER | HG003 | 98.76% | 94.14% | 96.39% | 99.66% | 99.74% | 99.70% | 89.64% | 57.16% | 69.80% | 88.38% | 60.95% | 72.15% | 91.02% | 53.61% | 67.48% | | | |
| | PEPPER | HG004 | 98.84% | 93.98% | 96.35% | 99.76% | 99.79% | 99.78% | 89.51% | 56.14% | 69.00% | 86.81% | 59.66% | 70.72% | 92.49% | 52.90% | 67.31% | | | |
| 80x | PEPPER | HG002 | 98.93% | 93.78% | 96.28% | 99.76% | 99.78% | 99.77% | 90.52% | 55.73% | 68.99% | 90.16% | 58.27% | 70.79% | 90.89% | 53.37% | 67.25% | 20463 | 35 | 349 |
| | PEPPER | HG003 | 98.85% | 94.19% | 96.46% | 99.66% | 99.75% | 99.71% | 90.54% | 57.47% | 70.31% | 89.00% | 61.40% | 72.67% | 92.25% | 53.79% | 67.96% | | | |
| | PEPPER | HG004 | 98.91% | 93.98% | 96.38% | 99.79% | 99.79% | 99.79% | 89.93% | 56.12% | 69.11% | 87.42% | 59.83% | 71.04% | 92.72% | 52.71% | 67.21% | | | |
| 90x | PEPPER | HG002 | 98.98% | 93.82% | 96.33% | 99.77% | 99.77% | 99.77% | 91.06% | 56.16% | 69.47% | 90.63% | 58.64% | 71.21% | 91.50% | 53.85% | 67.80% | 20373 | 35 | 321 |
| | PEPPER | HG003 | 98.84% | 94.26% | 96.50% | 99.67% | 99.75% | 99.71% | 90.55% | 58.00% | 70.71% | 89.23% | 61.91% | 73.10% | 91.99% | 54.36% | 68.33% | | | |
| | PEPPER | HG004 | 98.90% | 94.06% | 96.42% | 99.79% | 99.82% | 99.80% | 89.97% | 56.57% | 69.46% | 87.24% | 60.14% | 71.19% | 93.00% | 53.29% | 67.75% | | | |
| 10x | Clair3-Trio | HG002 | 96.71% | 89.30% | 92.85% | 98.07% | 96.55% | 97.30% | 81.05% | 43.34% | 56.48% | 80.65% | 44.05% | 56.98% | 81.44% | 42.67% | 56.00% | 7072 | 24 | 494 |
| | Clair3-Trio | HG003 | 94.92% | 89.48% | 92.12% | 96.44% | 96.55% | 96.49% | 77.07% | 42.85% | 55.07% | 78.75% | 44.68% | 57.01% | 75.44% | 41.13% | 53.24% | | | |
| | Clair3-Trio | HG004 | 94.63% | 89.15% | 91.81% | 95.96% | 96.41% | 96.18% | 78.50% | 41.83% | 54.58% | 80.17% | 43.19% | 56.13% | 76.92% | 40.59% | 53.14% | | | |
| 20x | Clair3-Trio | HG002 | 98.45% | 93.51% | 95.91% | 99.59% | 99.49% | 99.54% | 87.30% | 55.57% | 67.91% | 86.90% | 56.41% | 68.41% | 87.69% | 54.79% | 67.44% | 6243 | 24 | 188 |
| | Clair3-Trio | HG003 | 98.23% | 93.46% | 95.79% | 99.39% | 99.42% | 99.40% | 86.30% | 54.13% | 66.53% | 86.70% | 55.46% | 67.64% | 85.91% | 52.88% | 65.47% | | | |
| | Clair3-Trio | HG004 | 98.39% | 93.35% | 95.81% | 99.46% | 99.50% | 99.48% | 87.18% | 53.33% | 66.18% | 87.22% | 54.42% | 67.02% | 87.15% | 52.33% | 65.39% | | | |
| 30x | Clair3-Trio | HG002 | 98.66% | 94.34% | 96.45% | 99.73% | 99.69% | 99.71% | 88.90% | 60.43% | 71.95% | 88.29% | 61.38% | 72.41% | 89.51% | 59.55% | 71.52% | 6935 | 31 | 176 |
| | Clair3-Trio | HG003 | 98.66% | 94.26% | 96.41% | 99.65% | 99.61% | 99.63% | 89.10% | 58.97% | 70.97% | 88.81% | 60.15% | 71.73% | 89.39% | 57.87% | 70.25% | | | |
| | Clair3-Trio | HG004 | 98.83% | 94.19% | 96.45% | 99.76% | 99.69% | 99.72% | 89.72% | 58.39% | 70.74% | 89.34% | 59.62% | 71.52% | 90.09% | 57.25% | 70.01% | | | |
| 40x | Clair3-Trio | HG002 | 98.73% | 94.84% | 96.75% | 99.76% | 99.74% | 99.75% | 89.79% | 63.79% | 74.59% | 89.16% | 64.59% | 74.91% | 90.41% | 63.04% | 74.29% | 7392 | 33 | 173 |
| | Clair3-Trio | HG003 | 98.82% | 94.86% | 96.80% | 99.69% | 99.70% | 99.69% | 90.82% | 62.91% | 74.33% | 90.69% | 64.52% | 75.40% | 90.94% | 61.42% | 73.32% | | | |
| | Clair3-Trio | HG004 | 98.83% | 94.72% | 96.73% | 99.77% | 99.75% | 99.76% | 90.11% | 61.90% | 73.39% | 90.21% | 63.36% | 74.44% | 90.01% | 60.56% | 72.41% | | | |
| 50x | Clair3-Trio | HG002 | 98.69% | 95.30% | 96.96% | 99.78% | 99.79% | 99.79% | 89.64% | 66.83% | 76.57% | 88.66% | 67.58% | 76.70% | 90.60% | 66.12% | 76.45% | 7748 | 33 | 215 |

| | | | | | | | | | | | | | | | | | | | |
|---|---|---|---|---|---|---|---|---|---|---|---|---|---|---|---|---|---|---|---|
| | Clair3-Trio | HG003 | 98.82% | 95.24% | 97.00% | 99.71% | 99.71% | 99.71% | 90.87% | 65.70% | 76.26% | 90.59% | 67.07% | 77.08% | 91.14% | 64.42% | 75.48% | | | |
| | Clair3-Trio | HG004 | 98.89% | 95.16% | 96.99% | 99.81% | 99.79% | 99.80% | 90.79% | 64.99% | 75.75% | 90.48% | 66.53% | 76.68% | 91.08% | 63.56% | 74.87% | | | |
| 60x | Clair3-Trio | HG002 | 98.72% | 95.54% | 97.10% | 99.78% | 99.75% | 99.77% | 90.14% | 68.84% | 78.07% | 89.31% | 69.80% | 78.36% | 90.96% | 67.96% | 77.79% | 8429 | 33 | 197 |
| | Clair3-Trio | HG003 | 98.81% | 95.47% | 97.11% | 99.72% | 99.72% | 99.72% | 90.90% | 67.39% | 77.40% | 90.75% | 69.25% | 78.56% | 91.06% | 65.65% | 76.30% | | | |
| | Clair3-Trio | HG004 | 98.90% | 95.43% | 97.13% | 99.81% | 99.79% | 99.80% | 91.14% | 66.98% | 77.22% | 90.85% | 68.09% | 77.84% | 91.41% | 65.97% | 76.63% | | | |
| 70x | Clair3-Trio | HG002 | 98.77% | 95.73% | 97.23% | 99.79% | 99.78% | 99.78% | 90.63% | 70.06% | 79.03% | 90.15% | 71.20% | 79.57% | 91.09% | 69.00% | 78.52% | 9052 | 32 | 183 |
| | Clair3-Trio | HG003 | 98.84% | 95.66% | 97.22% | 99.74% | 99.72% | 99.73% | 91.19% | 68.86% | 78.47% | 90.96% | 70.44% | 79.40% | 91.42% | 67.38% | 77.58% | | | |
| | Clair3-Trio | HG004 | 98.94% | 95.57% | 97.22% | 99.80% | 99.80% | 99.80% | 91.55% | 68.01% | 78.04% | 91.09% | 69.23% | 78.67% | 91.99% | 66.89% | 77.46% | | | |
| 80x | Clair3-Trio | HG002 | 98.79% | 95.85% | 97.29% | 99.79% | 99.81% | 99.80% | 90.79% | 70.75% | 79.53% | 90.16% | 71.72% | 79.89% | 91.41% | 69.86% | 79.19% | 9096 | 33 | 185 |
| | Clair3-Trio | HG003 | 98.86% | 95.77% | 97.29% | 99.72% | 99.75% | 99.73% | 91.64% | 69.55% | 79.08% | 91.33% | 71.06% | 79.93% | 91.94% | 68.13% | 78.26% | | | |
| | Clair3-Trio | HG004 | 99.04% | 95.70% | 97.34% | 99.81% | 99.80% | 99.81% | 92.47% | 68.95% | 78.99% | 92.28% | 70.24% | 79.76% | 92.65% | 67.77% | 78.28% | | | |
| 90x | Clair3-Trio | HG002 | 98.78% | 95.91% | 97.33% | 99.80% | 99.82% | 99.81% | 90.74% | 71.17% | 79.77% | 90.23% | 72.02% | 80.10% | 91.23% | 70.39% | 79.46% | 9076 | 32 | 198 |
| | Clair3-Trio | HG003 | 98.93% | 95.84% | 97.36% | 99.76% | 99.75% | 99.76% | 91.85% | 70.00% | 79.45% | 92.25% | 71.82% | 80.76% | 91.45% | 68.29% | 78.19% | | | |
| | Clair3-Trio | HG004 | 99.03% | 95.70% | 97.33% | 99.83% | 99.81% | 99.82% | 92.28% | 68.92% | 78.91% | 92.33% | 69.97% | 79.61% | 92.22% | 67.96% | 78.25% | | | |

Supplementary Table 2. HG002 trio's *de novo* variants failed to be detected by Clair3-Trio.

| CHR | POS | REF | ALT | found by Clair3 | found by PEPPER | found by Clair3-Trio |
|---|---|---|---|---|---|---|
| chr20 | 3390371 | CT | C | N | N | N |
| chr20 | 18487996 | C | G | Y | Y | N |
| chr20 | 36652410 | C | CA | N | N | N |
| chr20 | 43658577 | A | C | Y | Y | N |
| chr20 | 47138749 | CA | C | N | N | N |
| chr20 | 51068541 | T | TA | N | N | N |
| chr20 | 57665372 | C | TA | N | N | N |

Supplementary Table 3. HG002 trio multiple-coverage benchmarking results while child’s coverage fixed at 60x.

| Tool | Sample | Coverage | Overall | | | SNP | | | INDEL | | | Insertion | | | Deletion | | | # of MCV | *de novo* TP | *de novo* FP |
|---|---|---|---|---|---|---|---|---|---|---|---|---|---|---|---|---|---|---|---|---|
| | | | Precision | Recall | F1-Score | Precision | Recall | F1-Score | Precision | Recall | F1-Score | Precision | Recall | F1-Score | Precision | Recall | F1-Score | | | |
| Clair3-Trio | HG002 | 60x | 98.63% | 94.43% | 96.49% | 99.70% | 99.65% | 99.67% | 89.02% | 61.37% | 72.66% | 88.22% | 62.34% | 73.06% | 89.81% | 60.48% | 72.28% | 10389 | 33 | 972 |
| Clair3-Trio | HG003 | 10x | 95.67% | 90.19% | 92.85% | 96.98% | 96.72% | 96.85% | 81.05% | 47.07% | 59.55% | 81.12% | 48.87% | 60.99% | 80.98% | 45.39% | 58.17% | | | |
| Clair3-Trio | HG004 | 10x | 95.41% | 89.93% | 92.59% | 96.60% | 96.65% | 96.62% | 81.89% | 46.19% | 59.06% | 81.67% | 47.32% | 59.92% | 82.10% | 45.14% | 58.25% | | | |
| Clair3-Trio | HG002 | 60x | 98.64% | 94.94% | 96.75% | 99.72% | 99.74% | 99.73% | 89.38% | 64.48% | 74.91% | 88.79% | 65.40% | 75.32% | 89.95% | 63.63% | 74.53% | 7437 | 33 | 406 |
| Clair3-Trio | HG003 | 20x | 98.34% | 93.76% | 96.00% | 99.41% | 99.44% | 99.43% | 87.48% | 56.29% | 68.50% | 87.54% | 57.95% | 69.74% | 87.41% | 54.74% | 67.32% | | | |
| Clair3-Trio | HG004 | 20x | 98.48% | 93.70% | 96.03% | 99.53% | 99.52% | 99.52% | 87.95% | 55.84% | 68.31% | 87.48% | 57.02% | 69.04% | 88.41% | 54.75% | 67.63% | | | |
| Clair3-Trio | HG002 | 60x | 98.72% | 95.18% | 96.92% | 99.76% | 99.76% | 99.76% | 89.97% | 66.16% | 76.25% | 89.36% | 66.97% | 76.57% | 90.55% | 65.41% | 75.95% | 7921 | 33 | 315 |
| Clair3-Trio | HG003 | 30x | 98.71% | 94.47% | 96.54% | 99.66% | 99.62% | 99.64% | 89.59% | 60.53% | 72.25% | 89.21% | 62.12% | 73.24% | 89.97% | 59.05% | 71.30% | | | |
| Clair3-Trio | HG004 | 30x | 98.81% | 94.39% | 96.55% | 99.77% | 99.71% | 99.74% | 89.68% | 59.77% | 71.73% | 89.14% | 61.12% | 72.52% | 90.21% | 58.52% | 70.99% | | | |
| Clair3-Trio | HG002 | 60x | 98.74% | 95.34% | 97.01% | 99.77% | 99.78% | 99.78% | 90.19% | 67.17% | 77.00% | 89.46% | 68.14% | 77.36% | 90.91% | 66.28% | 76.66% | 7860 | 32 | 248 |
| Clair3-Trio | HG003 | 40x | 98.79% | 94.95% | 96.84% | 99.69% | 99.71% | 99.70% | 90.56% | 63.56% | 74.70% | 90.18% | 65.34% | 75.77% | 90.93% | 61.91% | 73.66% | | | |
| Clair3-Trio | HG004 | 40x | 98.82% | 94.82% | 96.78% | 99.79% | 99.75% | 99.77% | 89.97% | 62.67% | 73.88% | 90.02% | 64.20% | 74.95% | 89.93% | 61.26% | 72.88% | | | |
| Clair3-Trio | HG002 | 60x | 98.73% | 95.45% | 97.06% | 99.79% | 99.76% | 99.78% | 90.06% | 68.10% | 77.55% | 89.22% | 69.14% | 77.90% | 90.88% | 67.13% | 77.22% | 7869 | 33 | 206 |
| Clair3-Trio | HG003 | 50x | 98.80% | 95.24% | 96.99% | 99.71% | 99.70% | 99.70% | 90.69% | 65.84% | 76.29% | 90.22% | 67.40% | 77.16% | 91.17% | 64.38% | 75.47% | | | |
| Clair3-Trio | HG004 | 50x | 98.87% | 95.21% | 97.00% | 99.80% | 99.79% | 99.79% | 90.68% | 65.38% | 75.98% | 90.32% | 66.86% | 76.84% | 91.03% | 64.02% | 75.17% | | | |
| Clair3-Trio | HG002 | 60x | 98.72% | 95.54% | 97.10% | 99.78% | 99.75% | 99.77% | 90.14% | 68.84% | 78.07% | 89.31% | 69.80% | 78.36% | 90.96% | 67.96% | 77.79% | 8429 | 33 | 197 |
| Clair3-Trio | HG003 | 60x | 98.81% | 95.47% | 97.11% | 99.72% | 99.72% | 99.72% | 90.90% | 67.39% | 77.40% | 90.75% | 69.25% | 78.56% | 91.06% | 65.65% | 76.30% | | | |
| Clair3-Trio | HG004 | 60x | 98.90% | 95.43% | 97.13% | 99.81% | 99.79% | 99.80% | 91.14% | 66.98% | 77.22% | 90.85% | 68.09% | 77.84% | 91.41% | 65.97% | 76.63% | | | |
| Clair3 | HG002 | 60x | 98.51% | 94.58% | 96.50% | 99.78% | 99.74% | 99.76% | 87.41% | 61.86% | 72.45% | 88.53% | 61.86% | 72.83% | 86.39% | 61.86% | 72.10% | 43205 | 35 | 3312 |

| | | | | | | | | | | | | | | | | | | | | |
|---|---|---|---|---|---|---|---|---|---|---|---|---|---|---|---|---|---|---|---|---|
| Clair3 | HG003 | 10x | 87.95% | 85.63% | 86.77% | 89.59% | 92.96% | 91.24% | 67.83% | 37.24% | 48.08% | 70.41% | 39.19% | 50.35% | 65.34% | 35.42% | 45.94% | | | |
| Clair3 | HG004 | 10x | 86.98% | 84.91% | 85.93% | 88.70% | 92.40% | 90.51% | 66.02% | 36.14% | 46.71% | 66.12% | 37.47% | 47.83% | 65.92% | 34.92% | 45.66% | | | |
| Clair3 | HG002 | 60x | 98.51% | 94.58% | 96.50% | 99.78% | 99.74% | 99.76% | 87.41% | 61.86% | 72.45% | 88.53% | 61.86% | 72.83% | 86.39% | 61.86% | 72.10% | 33565 | 35 | 1029 |
| Clair3 | HG003 | 20x | 97.07% | 92.61% | 94.79% | 98.92% | 98.84% | 98.88% | 78.76% | 51.46% | 62.25% | 78.48% | 53.16% | 63.38% | 79.04% | 49.88% | 61.16% | | | |
| Clair3 | HG004 | 20x | 97.07% | 92.56% | 94.76% | 99.11% | 98.92% | 99.02% | 77.39% | 51.07% | 61.53% | 75.49% | 52.71% | 62.08% | 79.35% | 49.56% | 61.01% | | | |
| Clair3 | HG002 | 60x | 98.51% | 94.58% | 96.50% | 99.78% | 99.74% | 99.76% | 87.41% | 61.86% | 72.45% | 88.53% | 61.86% | 72.83% | 86.39% | 61.86% | 72.10% | 32460 | 35 | 680 |
| Clair3 | HG003 | 30x | 97.82% | 93.92% | 95.83% | 99.45% | 99.51% | 99.48% | 82.57% | 57.00% | 67.44% | 82.08% | 58.75% | 68.48% | 83.07% | 55.36% | 66.44% | | | |
| Clair3 | HG004 | 30x | 98.02% | 93.80% | 95.86% | 99.66% | 99.58% | 99.62% | 82.69% | 56.14% | 66.88% | 80.50% | 58.18% | 67.55% | 84.97% | 54.26% | 66.23% | | | |
| Clair3 | HG002 | 60x | 98.51% | 94.58% | 96.50% | 99.78% | 99.74% | 99.76% | 87.41% | 61.86% | 72.45% | 88.53% | 61.86% | 72.83% | 86.39% | 61.86% | 72.10% | 31502 | 35 | 575 |
| Clair3 | HG003 | 40x | 98.43% | 94.41% | 96.38% | 99.62% | 99.68% | 99.65% | 87.21% | 59.65% | 70.84% | 87.49% | 61.69% | 72.36% | 86.94% | 57.74% | 69.39% | | | |
| Clair3 | HG004 | 40x | 98.55% | 94.27% | 96.37% | 99.76% | 99.69% | 99.72% | 87.27% | 58.95% | 70.37% | 85.18% | 60.95% | 71.06% | 89.43% | 57.11% | 69.70% | | | |
| Clair3 | HG002 | 60x | 98.51% | 94.58% | 96.50% | 99.78% | 99.74% | 99.76% | 87.41% | 61.86% | 72.45% | 88.53% | 61.86% | 72.83% | 86.39% | 61.86% | 72.10% | 31465 | 35 | 501 |
| Clair3 | HG003 | 50x | 98.60% | 94.76% | 96.64% | 99.67% | 99.70% | 99.68% | 88.81% | 62.19% | 73.15% | 89.03% | 64.38% | 74.73% | 88.59% | 60.14% | 71.64% | | | |
| Clair3 | HG004 | 50x | 98.74% | 94.70% | 96.68% | 99.80% | 99.74% | 99.77% | 89.04% | 61.81% | 72.97% | 87.23% | 63.88% | 73.75% | 90.90% | 59.92% | 72.22% | | | |
| Clair3 | HG002 | 60x | 98.51% | 94.58% | 96.50% | 99.78% | 99.74% | 99.76% | 87.41% | 61.86% | 72.45% | 88.53% | 61.86% | 72.83% | 86.39% | 61.86% | 72.10% | 30725 | 35 | 458 |
| Clair3 | HG003 | 60x | 98.67% | 95.10% | 96.85% | 99.70% | 99.73% | 99.71% | 89.51% | 64.57% | 75.02% | 90.06% | 66.06% | 76.21% | 88.96% | 63.18% | 73.89% | | | |
| Clair3 | HG004 | 60x | 98.88% | 95.08% | 96.94% | 99.81% | 99.76% | 99.79% | 90.60% | 64.56% | 75.39% | 89.38% | 66.95% | 76.56% | 91.84% | 62.36% | 74.28% | | | |
| PEPPER | HG002 | 60x | 98.61% | 93.48% | 95.98% | 99.68% | 99.70% | 99.69% | 87.81% | 54.09% | 66.94% | 86.99% | 56.58% | 68.56% | 88.67% | 51.78% | 65.38% | 39971 | 35 | 3556 |
| PEPPER | HG003 | 10x | 54.01% | 83.86% | 65.70% | 54.89% | 91.51% | 68.62% | 41.90% | 33.37% | 37.15% | 37.34% | 37.35% | 37.35% | 48.92% | 29.65% | 36.92% | | | |
| PEPPER | HG004 | 10x | 50.55% | 83.40% | 62.95% | 51.21% | 91.22% | 65.60% | 40.95% | 32.50% | 36.24% | 35.56% | 36.33% | 35.94% | 49.50% | 28.97% | 36.55% | | | |
| PEPPER | HG002 | 60x | 98.61% | 93.48% | 95.98% | 99.68% | 99.70% | 99.69% | 87.81% | 54.09% | 66.94% | 86.99% | 56.58% | 68.56% | 88.67% | 51.78% | 65.38% | 25539 | 35 | 867 |
| PEPPER | HG003 | 20x | 87.96% | 92.13% | 90.00% | 89.30% | 98.97% | 93.89% | 73.03% | 46.95% | 57.15% | 71.07% | 51.42% | 59.67% | 75.35% | 42.77% | 54.56% | | | |
| PEPPER | HG004 | 20x | 87.86% | 91.92% | 89.84% | 89.28% | 99.00% | 93.89% | 71.98% | 45.77% | 55.96% | 67.84% | 49.75% | 57.41% | 77.07% | 42.11% | 54.46% | | | |
| PEPPER | HG002 | 60x | 98.61% | 93.48% | 95.98% | 99.68% | 99.70% | 99.69% | 87.81% | 54.09% | 66.94% | 86.99% | 56.58% | 68.56% | 88.67% | 51.78% | 65.38% | 23138 | 35 | 607 |
| PEPPER | HG003 | 30x | 96.97% | 93.24% | 95.07% | 98.52% | 99.54% | 99.03% | 81.08% | 51.65% | 63.10% | 78.89% | 55.90% | 65.44% | 83.63% | 47.68% | 60.73% | | | |
| PEPPER | HG004 | 30x | 97.14% | 93.07% | 95.06% | 98.74% | 99.60% | 99.17% | 80.69% | 50.50% | 62.12% | 76.23% | 54.35% | 63.45% | 86.04% | 46.96% | 60.76% | | | |
| PEPPER | HG002 | 60x | 98.61% | 93.48% | 95.98% | 99.68% | 99.70% | 99.69% | 87.81% | 54.09% | 66.94% | 86.99% | 56.58% | 68.56% | 88.67% | 51.78% | 65.38% | 21688 | 35 | 499 |
| PEPPER | HG003 | 40x | 98.05% | 93.64% | 95.79% | 99.38% | 99.66% | 99.52% | 84.53% | 53.90% | 65.83% | 82.31% | 58.18% | 68.17% | 87.08% | 49.90% | 63.45% | | | |
| PEPPER | HG004 | 40x | 98.16% | 93.47% | 95.76% | 99.56% | 99.70% | 99.63% | 84.02% | 52.90% | 64.92% | 80.30% | 56.80% | 66.53% | 88.37% | 49.31% | 63.30% | | | |
| PEPPER | HG002 | 60x | 98.61% | 93.48% | 95.98% | 99.68% | 99.70% | 99.69% | 87.81% | 54.09% | 66.94% | 86.99% | 56.58% | 68.56% | 88.67% | 51.78% | 65.38% | 21021 | 35 | 474 |
| PEPPER | HG003 | 50x | 98.41% | 93.86% | 96.08% | 99.57% | 99.69% | 99.63% | 86.66% | 55.40% | 67.59% | 84.83% | 59.53% | 69.96% | 88.71% | 51.54% | 65.20% | | | |
| PEPPER | HG004 | 50x | 98.50% | 93.70% | 96.04% | 99.68% | 99.73% | 99.70% | 86.54% | 54.39% | 66.80% | 82.91% | 58.30% | 68.46% | 90.74% | 50.79% | 65.13% | | | |
| PEPPER | HG002 | 60x | 98.61% | 93.48% | 95.98% | 99.68% | 99.70% | 99.69% | 87.81% | 54.09% | 66.94% | 86.99% | 56.58% | 68.56% | 88.67% | 51.78% | 65.38% | 20559 | 35 | 455 |
| PEPPER | HG003 | 60x | 98.64% | 94.02% | 96.27% | 99.61% | 99.72% | 99.67% | 88.70% | 56.35% | 68.91% | 86.80% | 60.31% | 71.17% | 90.82% | 52.65% | 66.66% | | | |
| PEPPER | HG004 | 60x | 98.74% | 93.89% | 96.26% | 99.76% | 99.77% | 99.77% | 88.37% | 55.61% | 68.27% | 85.10% | 59.24% | 69.86% | 92.07% | 52.28% | 66.69% | | | |

### Supplementary Notes

#### Summary of methods tested that showed no or negligible improvement

(1) Adding position channels for trios: We added position channels in the Trio-to-Trio model to indicate the sample role in trios, including the child, parent1 and parent2 position channels. However, we observed that adding the position channel slightly decreased (-0.1%) the performance at the training terminal, when trained at chromosome 1 and tested at chromosome 20. We think that Clair3-Trio input, which is position order specific, is already encoded with position information, so it doesn't need the position channels for trio variant calling.

(2) Training with all data, including the Mendelian inheritance violation (MCV) variants for Clair3-Trio models: We found marginal MCV variants in the GIAB true set (1442/4924197). We included the MCV sites in the training data and found no improvement in overall variant prediction accuracy, but higher predicted MCV (+382, tested at chromosome 20). Although including MCV variants enriched the training data, we found that including them may deter model training performance, as it results in a conflict with the MCVLoss function introduced in Clair3-Trio.

(3) Trio phasing: We phased alignment from trio based on the WhatsHap phasing with the pedigree option. We found that WhatsHap phasing with the pedigree option in ONT data results in a similar number of phased sites among different depths compared to individual phasing. Furthermore, extensive testing for trio phasing for calling variants in Clair3-Trio resulted in no significant difference compared to phasing alone. We believe that the individuals' phasing module for ONT data is robust enough at present to gather information for variant calling.

#### Data sources

##### Reference genomes

*GRCh38_no_alt*

https://ftp.ncbi.nlm.nih.gov/genomes/all/GCA/000/001/405/GCA_000001405.15_GRCh38/seqs_for_alignment_pipelines.ucsc_ids/GCA_000001405.15_GRCh38_no_alt_analysis_set.fna.gz

#### GIAB Truth Variants

*HG002 (NA24385), GRCh38, v4.2.1*

https://ftp-trace.ncbi.nlm.nih.gov/giab/ftp/release/AshkenazimTrio/HG002_NA24385_son/NISTv4.2.1/GRCh38/

*HG003 (NA24149), GRCh38, v4.2.1*

https://ftp-trace.ncbi.nlm.nih.gov/giab/ftp/release/AshkenazimTrio/HG003_NA24149_father/NISTv4.2.1/GRCh38/

*HG004 (NA24143), GRCh38, v4.2.1*

https://ftp-trace.ncbi.nlm.nih.gov/giab/ftp/release/AshkenazimTrio/HG004_NA24143_mother/NISTv4.2.1/GRCh38/

#### Oxford Nanopore (ONT) Sequencing Data

*HG002 HD (NA24385), GRCh38_no_alt, 432.38-fold*

https://s3-us-west-2.amazonaws.com/human-pangenomics/index.html?prefix=NHGRI_UCSC_panel/HG002/nanopore/Guppy_4.2.2/

*HG003 Guppy 4.2.2 (NA24149), GRCh38_no_alt, 84.97-fold*

https://s3-us-west-2.amazonaws.com/human-pangenomics/index.html?prefix=NHGRI_UCSC_panel/HG003/nanopore/Guppy_4.2.2/

*HG004 GUPPY4.2.2 (NA24143), GRCh38_no_alt, 87.51-fold*

https://s3-us-west-2.amazonaws.com/human-pangenomics/index.html?prefix=NHGRI_UCSC_panel/HG004/nanopore/Guppy_4.2.2/

#### Commands

##### Read alignment using Minimap2 (v2.17-r941)

```
# Align ONT reads to GRCh38_no_alt
minimap2 -t ${THREADS} -aL -z 600,200 -x map-ont ref.fa input.fastq.gz | samtools view -bh
-o output.unsorted.bam -
samtools sort -@ ${THREADS} -o output.sorted.bam output.unsorted.bam
samtools index -@ ${THREADS} output.sorted.bam
```

##### BAM subsampling using Samtools (v1.10)

```
# Using ${FRAC} for both random seed and subsampling fraction
samtools view -@ ${THREADS} -s ${FRAC}.${FRAC} -b -o subsampled.bam ${BAM}
samtools index -@ ${THREADS} subsampled.bam
```

##### Coverage calculation using Mosdepth (v0.2.9)

```
mosdepth -t ${THREADS} -n -x --quantize 0:15:150: output ${BAM}
```

##### Clair3-Trio model training

Training section at: https://github.com/HKU-BAL/Clair3-Trio.

##### Running Clair3-Trio (v0.1)

```
${CLAIR3_TRIO_DIR}/run_clair3_trio.sh \
  --bam_fn_c=${BAM_C} \
  --bam_fn_p1=${BAM_P1} \
  --bam_fn_p2=${BAM_P2} \
```

```
  --ref_fn=${REF} \
  --threads=${THREADS} \
  --model_path_clair3="${MODEL_DIR_C3}" \
  --model_path_clair3_trio="${MODEL_DIR_C3T}" \
  --trio_model_prefix="${TRIO_M_PREFIX}" \
  --output=${OUTPUT_DIR} \
  --sample_name_c=${SAMPLE_C} \
  --sample_name_p1=${SAMPLE_P1} \
  --sample_name_p2=${SAMPLE_P2} \
```

Running Clair3 (v0.1-r6)

```
docker run -it \
  -v ${INPUT_DIR}:${INPUT_DIR} \
  -v ${OUTPUT_DIR}:${OUTPUT_DIR} \
  hkubal/clair3: v0.1-r6\
  /opt/bin/run_clair3.sh \
  --bam_fn=${INPUT_DIR}/input.bam \
  --ref_fn=${INPUT_DIR}/ref.fa \
  --threads=${THREADS} \
  --platform="${PLATFORM}" \
  --model_path="/opt/models/${PLATFORM}" \
  --output=${OUTPUT_DIR}
```

Running PEPPER (r0.4)

```
docker run --ipc=host \
  -v "${INPUT_DIR}":"${INPUT_DIR}" \
  -v "${OUTPUT_DIR}":"${OUTPUT_DIR}" \
  kishwars/pepper_deepvariant:r0.4 \
  run_pepper_margin_deepvariant call_variant \
  -b "${BAM}" \
  -f "${REF}" \
```

```
  -o "${OUTPUT_DIR}" \
  -p "${SAMPLE}" \
  -t ${THREADS} \
  --ont
```

#### Benchmarking using hap.py (v0.3.12)

```
hap.py ${GIAB_BASELINE_VCF} output.vcf.gz \
  -o ${OUTPUT_DIR}/happy \
  -r ${REF} \
  -f ${GIAB_CONFIDENT_BED} \
  --threads ${THREADS} \
  --pass-only \
  --engine=vcfeval
```

#### Merge VCF with BCFtools (v1.12)

```
M_VCF = merged_TRIO.vcf.gz
M_VCF_annotated = merged_annotated_TRIO_ann.vcf.gz

${BCFTOOLS} merge child.vcf.gz \
 Parent1.vcf.gz \
 Parent2. vcf.gz \
 --threads 32 -f PASS -0 -m all| ${BCFTOOLS} view -O z -o ${M_VCF}
 ${BCFTOOLS} index ${M_VCF}
```

#### Benchmarking using RTG tools (v3.12.1)

```
cat $PED
       #fam-id ind-id pat-id mat-id sex phen
       1 HG002 HG003 HG004 1 0
       1 HG003 0 0 1 0
       1 HG004 0 0 2 0
```

```
${RTG} mendelian -i ${M_VCF} -o ${M_VCF_annotated} --pedigree ${PED} -t
${REF_SDF_FILE_PATH}
```

##### Computing the TP and FP of number of *de novo* variants

```
${BCFTOOLS} merge --threads 8 -f PASS -0 -m all ${GIAB Truth Variants HG002} ${GIAB Truth
Variants HG003} ${GIAB Truth Variants HG004} | ${BCFTOOLS} view -O z -o ${TRIO_GIAB_MERGED
}
 ${BCFTOOLS} index ${TRIO_GIAB_MERGED}
${CLAIR3_TRIO} Check_de_novo --call_vcf ${M_VCF} --ctgName chr20 --bed_fn PED --true_vcf
$TRIO_GIAB_MERGED
```
